# Claudin-12 deficiency causes nerve barrier breakdown, mechanical hypersensitivity and painfulness in polyneuropathy

**DOI:** 10.1101/768267

**Authors:** Jeremy Tsung-Chieh Chen, Xiawei Hu, Kathrin Doppler, Olga Breitkreuz-Korff, Isabel U. C. Otto, Joachim Schwabe, Ann-Kristin Reinhold, Dorothee Günzel, Sophie Dithmer, Mohammed K. Hankir, Petra Fallier-Becker, Lars Winkler, Rosel Blasig, Claudia Sommer, Alexander Brack, Ingolf E. Blasig, Heike L. Rittner

**Author notes:** These authors contributed equally to this work (Jeremy Tsung-Chieh Chen, Xiawei Hu). **Corresponding author:** Heike Rittner, Dept. Anesthesiology and Critical Care, University Hospital of Würzburg, Oberdürrbacher Strasse 6, D-97080 Würzburg, Germany.

## Abstract

Peripheral nerves and their axons are shielded by the blood-nerve and the myelin barrier, but understanding of how these barriers impact nociception is limited. Here, we identified a regulatory axis of the tight junction protein claudin-12, sex-dependently controlling perineurial and myelin barrier integrity. In nerve biopsies, claudin-12 in Schwann cells was lost in male and postmenopausal female patients with painful but not painless polyneuropathy. Global *Cldn12* gene-knockout selectively increased perineurial/myelin barrier leakage, damaged tight junction protein expression and morphology, increased proinflammatory cytokines and induced mechanical hypersensitivity in naïve and neuropathic male mice, respectively. Other barriers and neurological function remained intact. *In vitro* transfection studies documented claudin-12 plasma membrane localisation without interaction with other tight junction proteins or intrinsic sealing properties. Rather, claudin-12 had a regulatory tight junction protein function on the myelin barrier via the morphogen SHH *in vivo* in *Cldn12-KO* and after local siRNA knockdown. Fertile female mice were completely protected. Collectively, these studies reveal the critical role of claudin-12 maintaining the myelin barrier and highlight restoration of the claudin-12/SHH pathway as a potential target for painful neuropathy.

## Introduction

Barriers are critical to shield delicate structures in the body. In the PNS, the blood nerve barrier (BNB) consists of the perineurium and endoneurial vessels. It is an important physiological fence which serves to maintain stable environments for axons, Schwann cells, and other endoneurial cells (Greathouse *et al*., 2016; Reinhold and Rittner, 2017). A second barrier in the PNS is the myelin barrier found at the paranode and the mesaxon of Schwann cells. Barriers are shaped by several main protein families: tight junction proteins, adherence junction proteins, and cytoplasmic accessory proteins (Neal and Gasque, 2016). Among the tight junction protein family, claudins [e.g. claudin-1, claudin-5, claudin-19, occludin and zona occludens (ZO-1, TPJ1)] have been identified as relevant for nerve barriers (Reinhold *et al*., 2018; Reinhold *et al*., 2019): Claudin-1 protects the perineurial barrier (Hackel *et al*., 2012; Sauer *et al*., 2014) and claudin-5, the major tight junction protein in the blood brain barrier (BBB), serves as a neuroprotector in in the endoneurial vessels (Moreau *et al*., 2016). Much less examined is the myelin barrier in neuropathy. Myelin tight junction proteins such as claudin-19 are responsible for sensory and motor transmissions (Guo *et al*., 2014). Peripheral myelin protein (PMP22) is necessary for the myelin barrier and dysregulated in inheritable neuropathies (Guo *et al*., 2014).

Claudin-12 is an atypical member of the claudin family because it is one of the few claudins that does not possess a PDZ binding motif. It is expressed in the intestine, urinary bladder, BBB and BNB in the nervous system (Shimizu *et al*., 2008; Markov *et al*., 2010; Amasheh *et al*., 2011; Reinhold and Rittner, 2017). Not very much is known about its function, except its upregulation in some types of cancer (Yang *et al*., 2015) and involvement in paracellular Ca^2+^ absorption (Fujita *et al*., 2008). However, its role in the BNB, myelin barrier and nociception are completely unknown.

Chronic inflammatory demyelinating polyneuropathy (CIDP) is a heterogenous (auto)immune-mediated inflammatory disease. Patients suffer from weakness, numbness and tingling and, in a subgroup, from pain (Kieseier *et al*., 2018). In some CIDP patients, autoantibodies against paranodal proteins are detected (Doppler *et al*., 2016; Dawes *et al*., 2018). In nerve biopsies, destruction of myelin sheaths and higher expression of inflammatory mediators such as TNF-α and IL-10 are found. Previous small studies have observed increased endoneurial edema at disease onset (Üceyler *et al*., 2016) and decreased claudin-5 and ZO-1 expression in sural nerve biopsies of CIDP patients (Kanda *et al*., 2004). This raises the question to the extent of which other tight junction proteins are expressed in the BNB and the myelin barrier are altered in CIDP patients and, if so, whether their expression is associated with painfulness.

Here, we specifically detected a loss of claudin-12 in Schwann cells in male and female postmenopausal patients with painful CIDP or non-inflammatory PNP. Using global *Cldn12*-KO mice, we uncovered that *Cldn12* deficiency resulted in perineurial and myelin barrier breakdown, downregulation of certain other tight junction proteins and exaggerated mechanical allodynia in naïve and neuropathic male mice. Moreover, a significant reduction of the morphogen and barrier stabilizer SHH in *Cldn12*-KO mice could explain the tight junction protein disruption. Together, this study clarifies the role of claudin-12 as a regulatory tight junction protein specifically in the perineurial/myelin but not BBB sealing.

## Methods

A detailed description of the methods is found in the supplementary information.

### Patients and tissue donors

This study was approved by the Ethical committee of the University of Würzburg (238/17) and is registered at the German clinical trial register (DRKS00017731). Patients with a diagnosis of CIDP fulfilling the Inflammatory Neuropathy Cause and Treatment (INCAT) criteria (Hughes *et al*., 2008) who attended the Dept of Neurology of the University Hospital Würzburg were included in the study. Sural nerve biopsies were obtained as part of the routine workup. The resulting cohort comprised of 10/12 patients with CIDP and 10/12 control patients with non-inflammatory PNP, respectively. The diagnosis of neuropathy was based on patient history and neurological examination and included pain assessment, quantitative sensory testing and electrophysiological tests. In addition, the overall disability sum score (ODSS) was obtained. It is composed of an arm and leg disability scale with a total score ranging from 0 (“no signs of disability”) to 12 (“most severe disability score”) (Merkies *et al*., 2002). Demographics and clinical characteristics of the different cohorts are given in the **Supplemental Table 1-3**. Female patients were only included if postmenopausal. Sural nerve biopsies were routinely analyzed for fiber loss (low, medium, high) and type of damage (axonal, demyelinating, mixed).

### Animals

Mouse experiments were performed in accordance with ARRIVE guidelines and approved by the Institutional Animal Care and Utilization Committee of the Government of Unterfranken, Germany (REG 2-264). For *Cldn12*-KO and WT mice, chimera with *Cldn12* gene ablating deletion were first generated by transfer of the murine embryonic stem cell clone 13208A (VelociGene; http://www.velocigene.com/komp/detail/13208) into blastocysts from albino-C57BL/6J-mice (B6(Cg)-Tyrc-2J/J) as provided by the Transgenic Core Facility, Max Planck Institute of Molecular Cell Biology and Genetics, Dresden/Germany. Then, the *Cldn12*-KO line (C57BL/6-Cldn12tm1(KOMP)Vlcg/Ph) was back-crossed ten times with C57BL/6 mice. From mating of the resulting heterozygous mice, KO and WT mice were investigated and revealed normal body-, behavior- and reproduction properties. The strain (EM:11196) is now available at EMMA strain search – Infrafrontier GmbH, Neuherberg/Germany (https://www.infrafrontier.eu/search). The animals were kept in small groups using standard cages, on 12:12 h dark–light cycle and standard diet according to the German animal welfare law approved by the animal ethics committee of Berlin (G0030/13, LaGeSo).

### Chronic constriction injury (CCI) and nociceptive tests

We housed mice in a light and temperature-controlled room under specific pathogen free conditions. Mice had free access to food and water. Male and female *Cldn12*-KO and WT-C57BL/6 mice (8-10 weeks) were randomized to the treatment vs. control. Under deep anesthesia (2% isoflurane), we placed four loose silk ligatures (6/0) (with 1 mm spacing) around the SN at the level of the right mid-thigh in *Cldn12*-KO and WT mice (Chen *et al*., 2014; Reinhold *et al*., 2019). Ligatures were tied until they elicited a brief twitch in the respective hind limb. The muscle layer and the incision in the shaved skin layer were closed with suturing material and metal clips separately after the surgery. Mice were monitored and maintained in a controlled temperature (37 °C) until fully recovered from anesthesia. The experimenter was blinded to the genotype.

*Thermal nociceptive behavior* responses were assessed by the Hargreaves method (Chen *et al*., 2014; Sauer *et al*., 2017; Reinhold *et al*., 2019). *Static mechanical hypersensitivity* was measured using von Frey filaments (Ugo Basile SRL, Italy) by using 50% paw withdrawal threshold (PWT) method (Chen *et al*., 2014; Sauer *et al*., 2017; Reinhold *et al*., 2019).

### Freeze-fracture electron microscopy

Approximately 4-6 mm of the SN (proximal to the sciatic trifurcation) from naïve WT and *Cldn12*-KO was gently freed from the surrounding connective tissue and fixed with 2.5% glutaraldehyde in PBS with Ca^2+^/Mg^2+^ for 2 h at RT (Wolburg *et al*., 2003). Quantification of TJs was conducted directly from the original images taken from 3 representative nerve preparations deriving from 4 individual animals in each group.

### RNAScope method

Cryosections of 10 µm thickness were cut at −20°C in a cryostat (Leica Biosystems CM3050 S Research Cryostat, Leica Biosystems Nussloch GmbH, Nussloch, Germany). Tissue sections were put in precooled 4% PFA in DEPC-treated distilled water. After 15 min of incubation at 4°C, samples were dehydrated in ethanol at RT. Hydrophobic barriers were drawn around the tissue sections. Afterwards, each section was incubated with two drops of RNAscope Protease IV reagent (15 min, RT). The RNAscope assay (Advanced Cell Diagnostics, Bio-Techne, Minneapolis, USA; 320850) was performed according to manufacturer’s instructions. All probes are found in the **Supplemental Table 4.** After blockage with 10% donkey serum in PBS, sections were immunostained with anti-claudin-1 Ab and secondary Ab to mark the perineurium. Finally, sections were mounted using VECTASHIELD Hardset Antifade Mounting Medium (H-1400, Vector Laboratories, Burlingame, USA).

All stainings (RNAscope, nerve permeability) were viewed using a KEYENCE BZ-9000 immunofluorescence microscope (Osaka, Japan) fitted with BZ-II analyzer software. The acquired images were analyzed using a free software package (ImageJ/Fiji). To analyze the fluorescence intensity of the immunostaining, the background was first measured in every picture. Using this corrected total cell fluorescence (CTCF) method, we calculated the integrated density, area and mean gray values of the fluorescence signal from red and green channels separately in each on region of interest (ROI).

### Nerve permeability

*Permeability of the BNB (perineurium):* Evans blue albumin (EBA) was prepared in 5% BSA with 1% EB dye (Sigma-Aldrich, St. Louis, USA) in sterile distilled PBS and filtered through a 0.20 µm filter (Sartorius stedim Biotech GmbH, Germany) (Hackel *et al*., 2012; Reinhold *et al*., 2019). Dissected SNs (1 cm length, proximal to the sciatic trifurcation) from *Cldn12*-KO and WT were *ex vivo* immersed in 2 ml of EBA for 1 h and then fixed with PFA (4% paraformaldehyde) overnight. Afterwards, tissues were embedded in Tissue–Tek, cut into 10 µm thick sections and mounted on microscope cover glasses with permaFluor Mountant (Thermo Scientific, Fremont, USA). The permeability of the perineurium was determined by the diffusion of EBA into the endoneurium of the SN.

*Permeability of the myelin barrier:* The whole SN was dissected into epiperineurium (EPN) and the desheathed SN (dSN). The dSNs were sealed with vaseline in both ends and incubated in artificial cerebrospinal fluid (as following (mM): 10 HEPES, 17.8 NaCl, 2 NaHCO3, 4 MgS04, 3.9 KCl, 3 KH2PO4, 1.2 CaCl2, 10 Dextrose; pH 7.4), containing fluorescein isothiocyanate-dextran (FD; 5 mg/ml; MW: 70 kDa; Sigma) for 1 h at 37 °C. Afterwards, nerves were fixed in 4% PFA for 5 min at RT, then placed on glass slides and teased into individual fibers under a dissecting microscope. We then mounted these teased fibers by VECTASHIELD Antifade Mounting Medium and covered them with microscope coverslips. The green fluorescence signal was determined.

*Permeability of the BNB (endoneurial vessels):* Anaesthetized mice were laid down in a supine position on a pad. The 5th intercostal space was opened and enlarged by a retractor to open the thorax. The pericardium was stripped exposing the heart anterior wall and 50 µl of FD solution was injected into the left ventricle using an insulin syringe. The dye-injected mice were sacrificed by decapitation after 2 min. SNs were dissected and embedded in Tissue-Tek. Frozen samples were cut into 12 µm-thick sections on a cryostat at −20°C. Without any fixation, microscope glass slides containing tissue sections were mounted and were imaged by fluorescence microscopy.

### Brain microvessel isolation and qRT-PCR

Brain capillaries were isolated from cortices of 8-20 weeks-old *Cldn12*-KO and WT mice by homogenization in Dulbecco’s modified Eagle’s medium (DMEM, 4.5 g/l glucose; Life Technologies, Darmstadt, Germany) with a dounce tissue grinder (Wheaton, Millville, USA) on ice. Myelin was removed by adding dextran (60-70 kDa, 16% (w/v) final concentration; Sigma-Aldrich) and centrifugation (15 min, 4,500×g, 4 °C). The resuspended pellet was filtered (40 µm nylon mesh, Merck Millipore) and the remaining capillaries were rinsed off with DMEM and 1% (v/v) fetal calf serum (Life Technologies) (Berndt *et al*., 2019).

RNA was extracted from isolated murine brain capillaries by GeneMATRIX Universal RNA Purification Kit (EURx, Gdansk, Poland). The cDNA was synthesized with the Maxima First Strand cDNA Syntheses Kit (ThermoFisher). qRT-PCR was performed with StepOne Real-Time PCR system, 48/96 well (Life Technologies, Darmstadt, Germany) using the Luminaris Color HiGreen high ROX qPCR Master Mix (ThermoFisher) as described in the manual. Primer sequences with a melting temperature of around 60 °C (BioTeZ, Berlin, Germany) designed with Primer3Plus.com (**Supplemental Table 5**). The mRNA expression was normalized to β-actin and displayed as fold change expression calculated by the 2^−ΔΔCT^ method as described above (Dithmer *et al*., 2017).

### Cell culture

Cell lines from human embryonic kidney (HEK-293; (Piontek *et al*., 2011)) and Madin-Darby canine kidney (MDCK-II) were grown in 6-well plates until 80% confluency and transfected using polyethylenimine (PEI; Polysciences Inc, Warrington, USA) (Cording *et al*., 2013). Constructs used were N-terminally fused with yellow fluorescent protein (YFP) or pmTurquoise2 (TRQ): YFP-*Cldn5* (Gehne *et al*., 2017), YFP-*Cldn12*, TRQ-*Cldn12* and YFP-*Ocln* (Bellmann *et al*., 2014). For each well, 10 µl of PEI (1 mg/ml) was added to 250 µl Opti-MEM (ThermoFisher); separately, 2 µg plasmid DNA was mixed with another 250 µl Opti-MEM. 5 min after incubation at RT, both solutions were combined and carefully mixed. Following 25 min incubation at RT, the mixture was added drop wise to each well. Transfection medium was removed after incubation overnight and cells were transferred into a 25 cm^2^ flask. Cells were cultured in DMEM (1 g/ml glucose) containing 10% fetal calf serum and 1% penicillin/streptomycin (Thermo Fischer) at 37 °C and 10% CO2. For transfected cells, G418 was added to the medium (Dithmer *et al*., 2017).

### Live cell imaging, trans-interaction, and transepithelial electrical resistance (TER)

Cells were seeded onto glass coverslips coated with poly-L-lysine (Sigma-Aldrich), washed with HBSS+/+ (with Ca^2+^ and Mg^2+^; ThermoFisher) and imaged in HBSS+/+ using a confocal microscope (Zeiss NLO with a Plan-Neofluar 100x 1.3 Oil objective, Zeiss, Oberkochen, Germany) (Cording *et al*., 2013). Plasma membranes were visualized by trypan blue (0.05%; Sigma-Aldrich) in HBSS+/+. For *trans*-interactions, the enrichment factor EF was detected. Half the fluorescence intensity (I) of a YFP- or TRQ-tagged protein at cell contacts between two transfected cells was divided by YFP or TRQ fluorescence at contacts between expressing and non-expressing cells (EF=Icontact/2Ino contact) (Piontek *et al*., 2008).

To measure the TER with a Cellzscope (nanoAnalytics, Muenster, Germany), cells were seeded onto hanging PET cell culture inserts (Merck Millipore) at a density of 80,000 cells/insert. After TER values had reached a stable plateau, the cells were transferred into a 24-well plate and washed twice with HBSS+/+ warmed to 37 °C (Staat *et al*., 2015).

### FRET measurements

HEK 293 cells (CRL-1573, A.T.C.C., Manassas, VA, USA; grown in Minimum Essential Medium Eagle (MEM; Sigma M0446) supplemented with 10% (v/v) FBS (Gibco 10270106) and 1% penicillin/streptomycin (Corning 30-002-CI) were transiently transfected with pcDNA 3.1 plasmids containing cDNA for N-terminally enhanced yellow or cyan fluorescent protein (EYFP or ECFP)-tagged murine *Cldn12*, human *CLDN19a* (NP_683763), human *CLDN19b* (NP_001116867) or human *PMP22* (NP_000295), respectively. To this end, HEK 293 cells were seeded onto poly-lysine (Sigma P48323) coated 32 mm diameter cover slips (Thermo Scientific/Menzel) in six well plates (Falcon 353046). On the following day, cells were co-transfected with two plasmids encoding a YFP- and a CFP-fusion protein, respectively. The total sum of plasmid DNA amounted to 4 µg/well but YFP/CFP ratios varied. DNA was mixed with 10 µl PEI (polyethylenimine, 1 mg/ml, Polysciences Europe GmbH, Eppelheim, Germany) in 500 µl MEM without supplements. After 30 min incubation, the mixture was added to the cells. They were used for live cell imaging in a Zeiss LSM 780 after 24 to 48 h. CFP and YFP fluorescence signals were excited at 458 nm (detection 463 – 507 nm) and 514 nm (detection 518 – 588 nm), respectively. FRET was determined as percent increase in CFP signal intensity before and after acceptor bleaching with 20 pulses of 514 nm laser radiation at 100% laser power (Milatz *et al*., 2015).

### Organ permeability

Male mice were i.v. injected with 0.5 mol/kg b.w. Na-fluorescein (NF; Sigma-Aldrich, Munich, Germany) or 0.26 mol/kg b.w. Evans blue (EB; Sigma-Aldrich), dissolved in saline. NF, 376 Da, served as small and EBA (due to its association to plasma albumin, 68 kDa) as large molecular weight markers. After 10 min, mice were anesthetized i.p. using ketamine (0.18 mg/g b.w.; CP-Pharma, Burgdorf, Germany) and xylazine (0.024 mg/g b.w.; Ceva Tiergesundheit, Düsseldorf, Germany), and perfused with 25 ml Dulbecco’s phosphate-buffered saline with Ca^2+^/Mg^2+^ (DPBS+/+; ThermoFisher Scientific, Darmstadt, Germany) containing 6250 IU heparin (Ratiopharm, Ulm, Germany). Organs were homogenized in DPBS+/+ followed by precipitation with trichloroacetic acid (Roth, Karlsruhe, Germany; final concentration 30% (w/v)) in the dark overnight at 4 °C. After centrifugation (19,000xg, 15 min), fluorescence of the supernatant was measured (NF 485/520 nm; EB 620/680 nm) with a plate reader (Tecan Safire; Tecan, Männedorf, Switzerland), and referred to organ weight (µgdye/mgorgan; (Dithmer *et al*., 2017)).

### siRNA treatment via perineurial injection

We use commercial siRNA to suppress local mRNA expression. *Cldn5*-targeting siRNA (Silencer® Pre-designed siRNA, siRNA ID: s64050, Ambion), *Cldn12*-targeting siRNA (Silencer® Pre-designed siRNA, siRNA ID: s82332, Ambion), *Cldn19*-targeting siRNA (Silencer® Pre-designed siRNA, siRNA ID: s202784, Ambion) or scrambled-siRNA-Cy3 (Mission®siRNA universal negative controls, ProducNo. SIC003, Sigma) alone were applied on the surface of the SN (2 µg of the target siRNA or scrambled-siRNA mixed with i-Fect™ (Neuromics, Edina, MN) in a ratio of 1:5 (W:V) to a final concentration of 400 mg/L).

### Fluorescence quantification

ImageJ/Fiji was used to analyze our targeting fluorescence signals. To measure FD signals in teased fibers, we scanned and imaged nerve fibers from *Cldn12*-KO and WT mice by fluorescence microscopy at 40X magnification. Microscope and camera settings (e.g., light level, exposure, gain, etc.) were identical for all images. The number of individual fibers with positive FD penetration and the total number of individual fibers were counted and calculated. The total number of fibers was approximately 100 in 5 mice from WT and KO groups. After background subtraction in every picture, bright field images were used to guide manual selection of individual nerve fiber outlines. 20 µm^2^ ROI were saved onto ROI manager. ROIs were applied on the green channel, and the selected FD signals were analyzed using the built-in plugin of ImageJ/Fiji. The mean gray values, area and integrated density of ROIs were obtained and analyzed. When the mean gray value of the ROI was bigger than 10, the related fiber was considered to be a positive FD penetration fiber. Finally, a ratio was calculated by dividing the number of positive FD penetration fibers by the total number of individual fibers.

To analyze claudin-12-immunoreactivity (IR) signals in the SN of human and mice, co-expression regions of S100 and claudin-12 staining were selected and taken from coronal sections of the SN. S100-IR signals were used to determine location of Schwann cells, which were saved to the ROI manager. These ROIs were applied in the red channel and claudin-12-IR signals in these ROIs were analyzed using the Measure built-in plugin of ImageJ/Fiji.

### Statistical analyses

Statistical analysis was done using SigmaPlot or Excel software. When comparisons were more than two groups, one- or two-way ANOVA followed by the Bonferroni post-hoc test was used. For pain behavior tests, two-way RM ANOVA with Bonferroni post-hoc test was employed. The two-tailed Student’s t-test was applied to examine differences between two groups. For transfected cell culture dataset, comparisons were performed using the Kruskal-Wallis-test and Dunns multiple comparison post hoc test. Differences between groups were considered significant if p □ 0.05. All data are expressed as mean ± SEM.

## Results

### Deficiency of claudin-12 in sural nerves of male or female postmenopausal patients with painful CIDP or non-inflammatory PNP

We investigated a panel of tight junction protein typical for each barrier in sural nerves in cohorts of male or female postmenopausal CIDP patients and patients with non-inflammatory idiopathic PNP **(Supplemental Table 1**). Claudin-1-immunoreactivity (IR) was detected in several layers of the perineurium (**Supplemental Fig S1A**). Both ZO-1-IR and occludin-IR were found in all compartments of the sural nerve, e.g. perineurium, Schwann cells and endoneurial vessels (**Supplemental Fig S1B, D**). Claudin-5-IR was spotted in the endoneurium in endoneurial blood vessels (**Supplemental Fig S1C**). Claudin-19-IR was observed in the perineurium and around nerve fibers co-localizing with the Schwann cell marker S100b (**Supplemental Fig S1E**). Claudin-1, -19 (in S100b positive cells) and ZO-1-IR were significantly reduced in PNP with severe fiber loss independently of inflammation or pain (**Supplemental Fig S1F-H**). No differences were found in the clinical characteristics of these groups. However, significantly more patients with CIDP were in the severe fiber loss group (**Supplemental Table 2**). Claudin-12-IR was predominantly located around nerve fibers of CIDP and non-inflammatory PNP patients and was co-expressed with S100b (**Fig. 1**). Some claudin-12-IR was observed in the perineurium. Patients with CIDP and non-inflammatory PNP with pain showed a significant reduction of claudin-12-IR in sural nerve samples compared to patients without pain (**Fig 1A, C**) unrelated to fiber loss and inflammation (**Fig 1C**). No differences were found in the clinical characteristics of these groups except for sex distribution (**Supplemental Table 3**). In summary, claudin-12 is unique because its loss parallels pain symptoms independently of fiber loss, inflammation, demyelinating/axonal neuropathy, functional impairment and disease duration.

**Figure 1.**
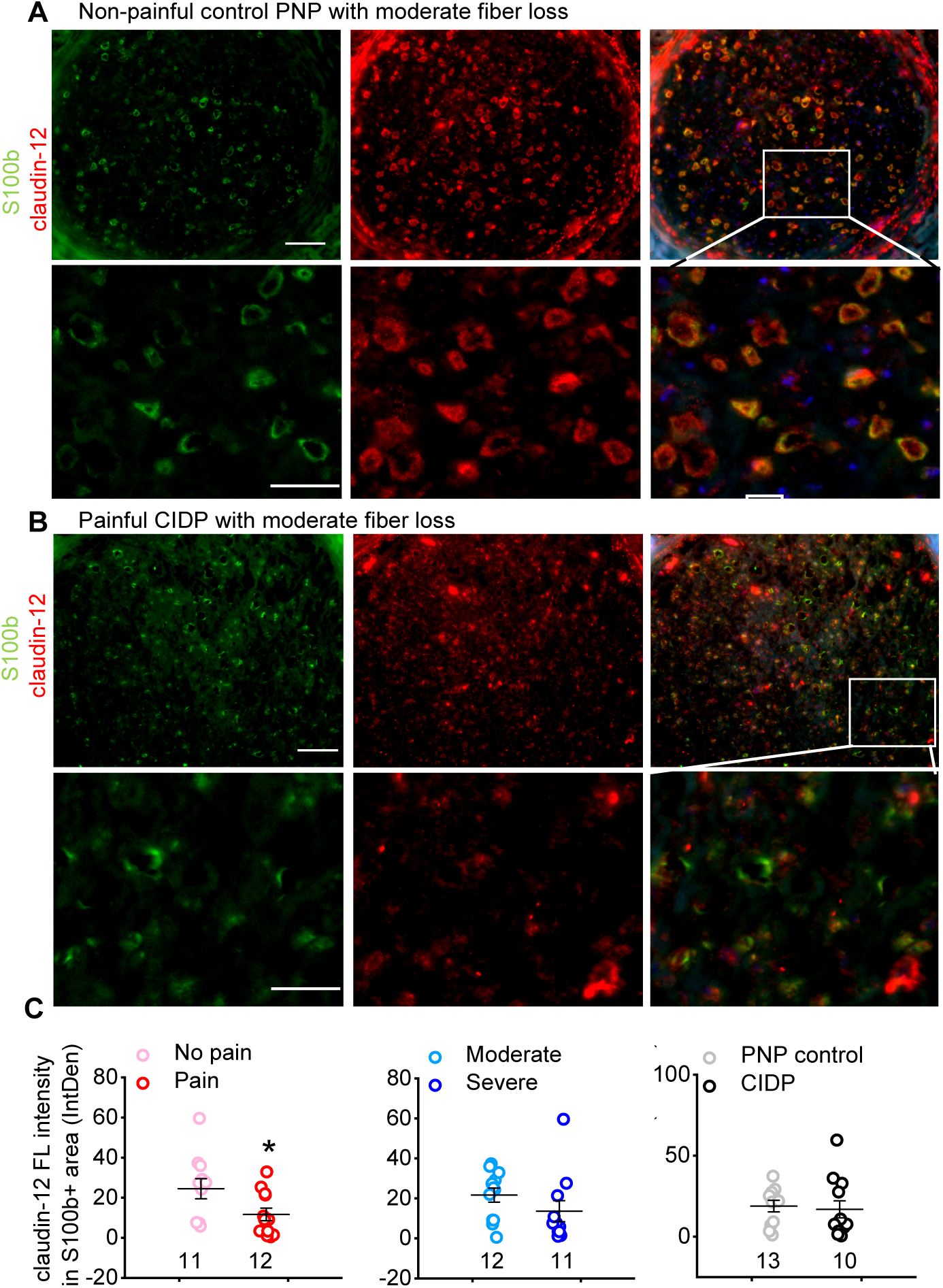
Reduced claudin-12-immunoreactivtiy (IR) in Schwann cells in sural nerves in a subgroup of patients with painful chronic inflammatory demyelinating polyneuropathy (CIDP) or non-inflammatory polyneuropathy (PNP). (**A-C**) Sections from sural nerves of 23 patients with either non-inflammatory PNP or CIDP were immunolabeled. Patients were grouped according to pain symptoms (red) and extend of fiber loss (blue) and presence/absence of inflammation (grey) (patients characteristics’: **Supplemental Table 2 and 3**). Representative immunostainings for claudin-12 (red) co-stained with the Schwann cell marker S100b (green) are displayed comparing a patient with (**A**) and without (**B**) pain and moderate fiber loss. Higher magnification merged views (white square) are shown in the lower row (scale bar upper row = 50 µm, scale bar lower low = 25 µm). (**C**) Claudin-12-IR within S100b+ was quantified according to patients’ subgroups with pain symptoms, extend of fiber loss and inflammation. PNP controls included idiopathic, vasculitic, hereditary and alcoholic PNP. The number of patients in each group are found below the dot plots. *p < 0.05, two-sided Student’s t-test. Data are shown as mean ± SEM.

### Claudin-12 and claudin-19 loss after nerve injury

Next, we chose to further examine the well-standardized mononeuropathy CCI in mice, characterized also by an inflammatory reaction due to the ligatures around the SN. Previous data from our and other labs revealed that CCI caused claudin-1, claudin-5, occludin and ZO-1 downregulation in the SN (Moreau *et al*., 2016; Reinhold *et al*., 2018; Reinhold *et al*., 2019). In WT mice, claudin-12-IR was found in myelinating, MBP-positive Schwann cells and S100b-positive Schwann cells (**Fig 2A**). Some claudin-12-IR was observed in the perineurium. After 7 d CCI, i.e. at the time point of mechanical hypersensitivity, *Cldn12* mRNA levels in the epiperineurium (EPN), desheathed sciatic nerve (dSN) and dorsal root ganglion (DRG) were significantly lower after CCI by 60%, 80% and 30 %, respectively (**Fig 2B**). Similarly, claudin-12-IR was decreased by 78.2% in the SN of male mice (**Fig 2C, D**). Since claudin-19 is mainly expressed in Schwann cells (Miyamoto *et al*., 2005; Guo *et al*., 2014), we then investigated claudin-19-IR in teased nerve fibers. Claudin-19-IR was significantly decreased in the paranode of injured mice by 50% (**Supplemental Fig S2A**). In summary, tight junction protein expression in Schwann cells is impaired in neuropathy.

**Figure 2.**
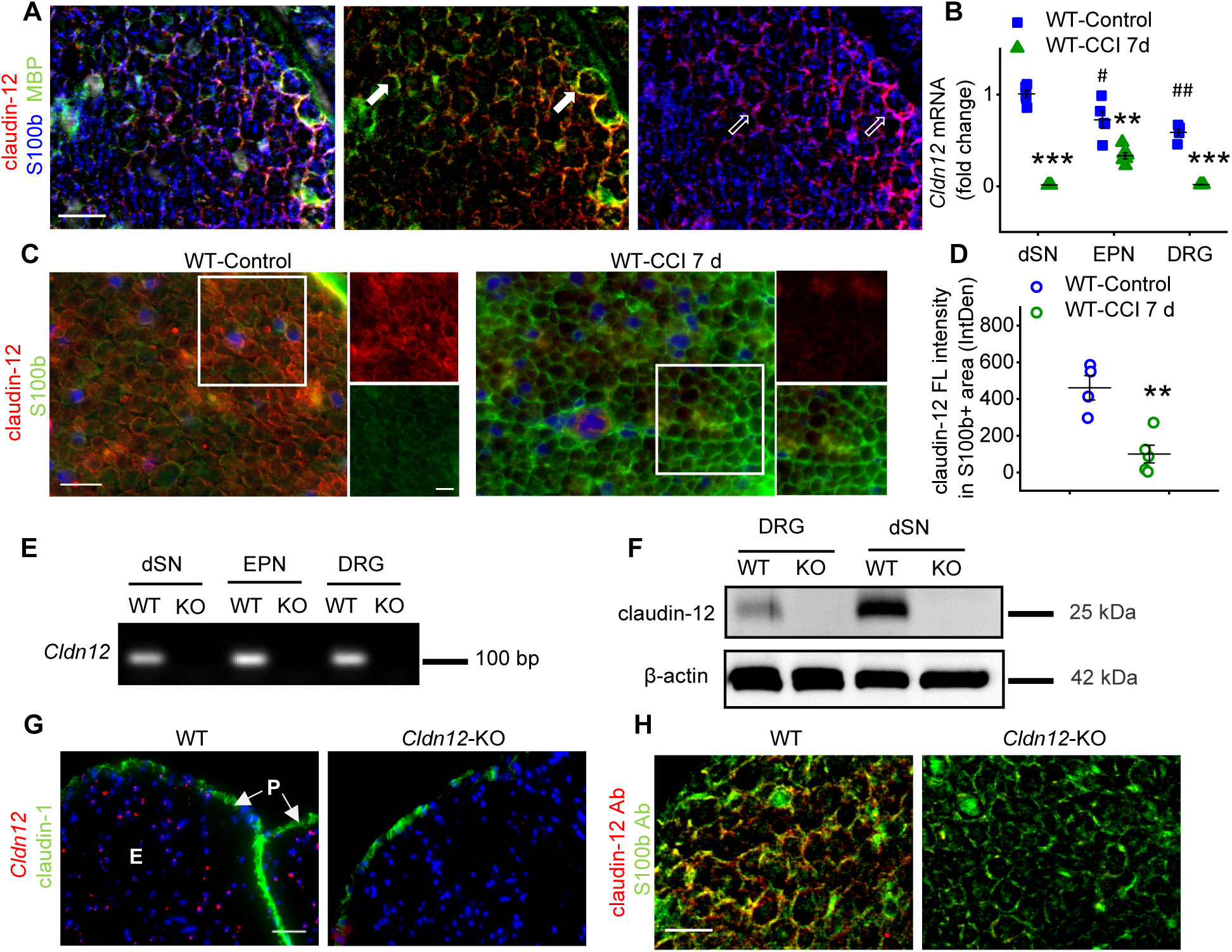
Claudin-12 loss in the sciatic nerve after nerve injury and generation of *Cldn12*-KO mice. WT mice underwent chronic constriction injury (CCI) for 7 d. (**A**) Triple labeling was performed with claudin-12 (red) and myelin basic protein (MBP; green) and S100b (blue) for Schwann cells in naïve mice [representative example; scale bar: 20 µm]. (**B**) *Cldn12* mRNA expression in the desheathed sciatic nerve (dSN), epiperineurium (EPN) and dorsal root ganglion (DRG) was analyzed in naïve and CCI mice. (##p < 0.01, EPN, DRG vs. dSN WT; **p < 0.01, ***p < 0.001, naïve versus CCI WT, two-way ANOVA followed by Bonferroni’s post hoc test; n = 6) (**C-D**) Double-label immunostaining was performed showing coexpression of claudin-12 (red) and S100b (green) in the sciatic nerve (SN) of untreated naive and CCI mice [representative example; scale bar image overview: 25 µm, scale bar magnified image (white square): 10 µm] and quantified (**p < 0.01, one-tailed Student’s t-test; n = 4-5). (**E, F**) *Cldn12*-KO mice were generated as described in the methods. Representative examples of *Cldn12* mRNA and protein expression in dSN, EPN and DRG tissues in *Cldn12*-KO and WT are displayed. Uncropped blots are provided in **Supplemental Fig S5A, B**. (**G**) Characteristic images of RNAscope® fluorescent assay for *Cldn12* mRNA (red) are presented which were co-stained with immunostaining for claudin-1 (green) and DAPI (blue) in the SN of WT and *Cldn12*-KO mice (scale bar: 50 µm). P: perineurium, E: endoneurium. (**H**) Immunofluorescence imaging of sectioned SN from WT and *Cldn12*-KO mice stained with claudin-12 (red) and S100b (green) antibody respectively (scale bar: 20 µm). Data are shown as mean ± SEM.

### Mechanical hypersensitivity in male *Cldn12*-KO mice

Because of the downregulated claudin-12-IR in male or female postmenopausal patients with painful CIDP/PNP and mice with nerve injury, we generated *Cldn12*-KO mice to investigate the functional role of claudin-12 in the BNB and myelin barrier. We confirmed the deletion of the *Cldn12* gene and protein in the PNS (**Fig 2E, F**) as well as several other tissues of KO mice (data not shown). To localize *Cldn12* mRNA expression in the PNS, we labelled SN sections with the *Cldn12* RNAscope probe and the anti-claudin-12 Ab (**Fig 2G, H**). *Cldn12* mRNA was detected in the endoneurium and perineurium lost in KO mice. The claudin-12-IR signal also was absent in the SN of *Cldn12*-KO mice. The summarized data confirmed the successful KO of *Cldn12*.

Male *Cldn12*-KO mice exhibited normal body weight (WT: 23.44 ± 0.71 g vs. *Cldn12*-KO: 22.13 ± 1.26 g) and normal physical appearance (**Supplemental Fig S3**). Gross neurological examination did not show any obvious sign of altered sensory or vestibular function in mutants. Specific evaluation of the auditory system using auditory brainstem responses, anxiety-related behavior (open field test, locomotor activity and rears), acoustic startle and pre-pulse inhibition, baseline level of immobility during habituation to the conditioning cages, contextual and cued freezing performance were also comparable between both genotypes. Only, the retinas of *Cldn12*-KO mice were slightly but significantly thinner than those of their WT littermates.

To determine whether the deletion of *Cldn12* is involved in motor and sensory function, we performed a battery of behavioral tests in *Cldn12-KO* and WT mice. Motor strength test for 2-paw and 4-paw handgrip as well as rotarod test showed no significant difference between KO and WT mice (**Fig 3A, B**). *Cldn12-*KO mice had significantly higher thresholds for electrical stimulation-elicited vocalization than WT mice, while thresholds for flinch and jump responses were comparable (**Fig 3C**). This indicates abnormal sensory processing after deletion of *Cldn12*. In the hot plate test, investigating supraspinal integrated thermal responses, no significant differences were observed (**Fig 3C**). However, male *Cldn12-*KO mice were more sensitive to mechanical stimuli (manual von Frey filaments test) at baseline levels and after CCI, while thermal sensation was unaffected (**Fig 3E, F**). Interestingly, no change in sensation of mechanical stimuli was observed in female *Cldn12*-KO, while both genotypes of fertile female mice still developed mechanical hypersensitivity after CCI (**Supplemental Fig S4A**). Together, these results demonstrate that *Cldn12* deficiency selectively affects mechanical sensation in male mice.

**Figure 3.**
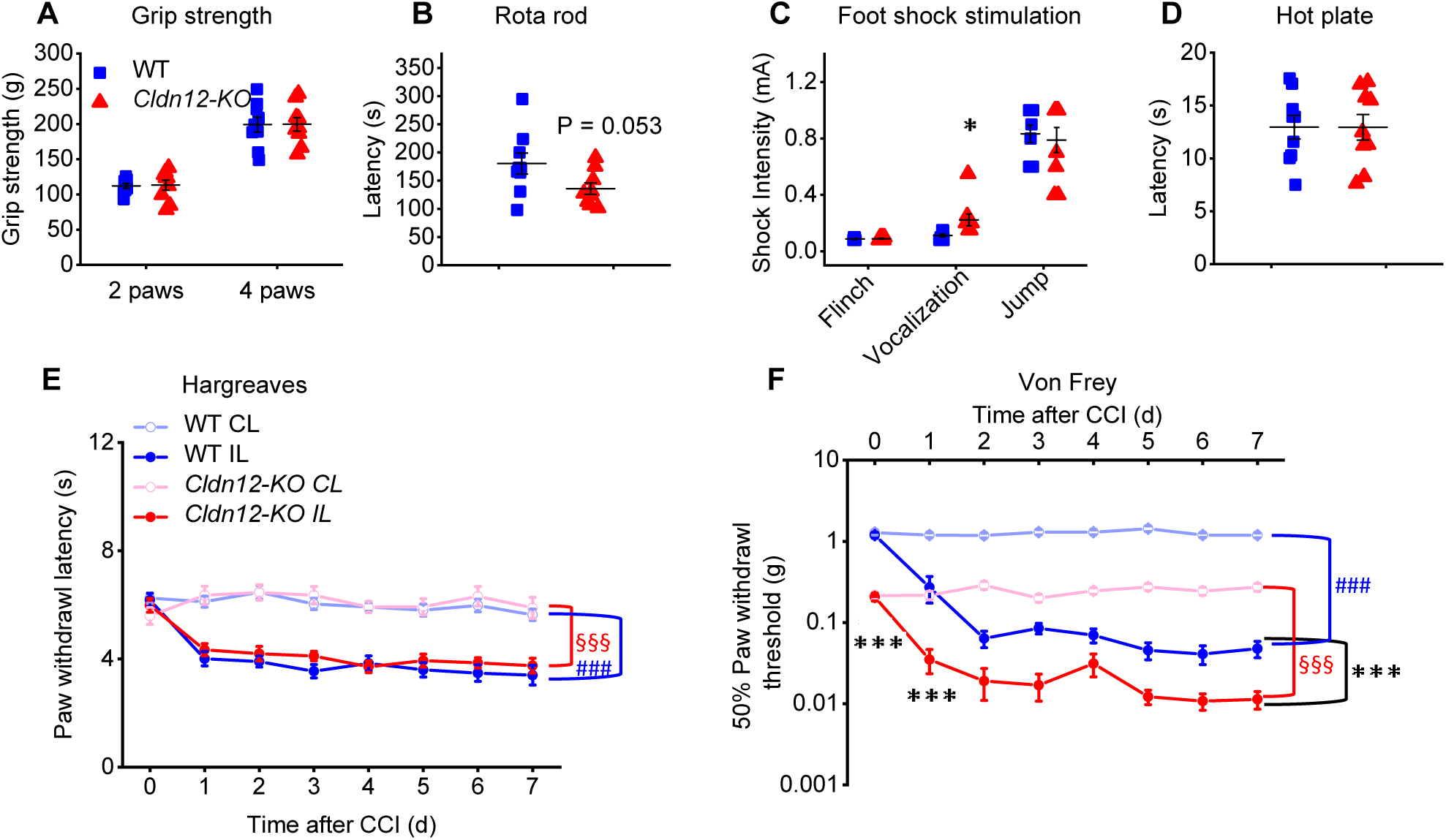
Mechanical hypersensitivity in male *Cldn12*-KO mice. (**A-D**) Behavioral tests were performed with naïve male *Cldn12*-KO mice in comparison with naïve male WT mice. (**A**) Slip thresholds of two and four paws were analyzed by the grip strength test. (**B**) The latency to fall of the Rota rod was tested in both genotypes. (**C**) Vocalization, flinch and jump thresholds to foot shock stimuli were obtained and averaged. (**D**) Nociceptive response latencies to a hot-plate surface were measured (all above *p < 0.05, unpaired two-sided Student’s t-test, n = 9). (**E, F**) Mice underwent CCI surgery. Nociceptive thresholds were obtained before and every day after CCI: Thermal hypersensitivity was detected in the Hargreaves test (**E**) and static mechanical hypersensitivity by Frey filaments test (**F**) (n = 10, IL: ipsilateral, CL: contralateral, two-way RM-ANOVA, Bonferroni’s post hoc test. ***p < 0.001 IL *Cldn12*-KO versus IL WT in the von Frey test only. §§§ p < 0.001 IL versus CL *Cldn12*-KO, ### p < 0.001 IL versus CL WT). Data are shown as mean ± SEM.

### Leakiness of the myelin barrier and the perineurium of the BNB in male *Cldn12*-KO mice

We next explored the morphological organization of the SN in *Cldn12*-KO (**Fig 4A**). Relative myelin thickness (G-ratio) and the number of undulated fibers were normal (**Fig 4B, C**). Undulated fibers were defined as fibers with >4 undulations on myelin (Yuan *et al*., 2018). The total number of axons was significantly lower in male *Cldn12*-KO (**Fig 4D**). Analyzing the distribution of axon and nerve fiber diameters, we found the increase in the percentage of axons with 7-9 µm diameter and a decrease in those with 3–7 µm diameter. In summary, the results could point towards a loss of small-diameter axons during embryonal development in *Cldn12*-KO.

**Figure 4.**
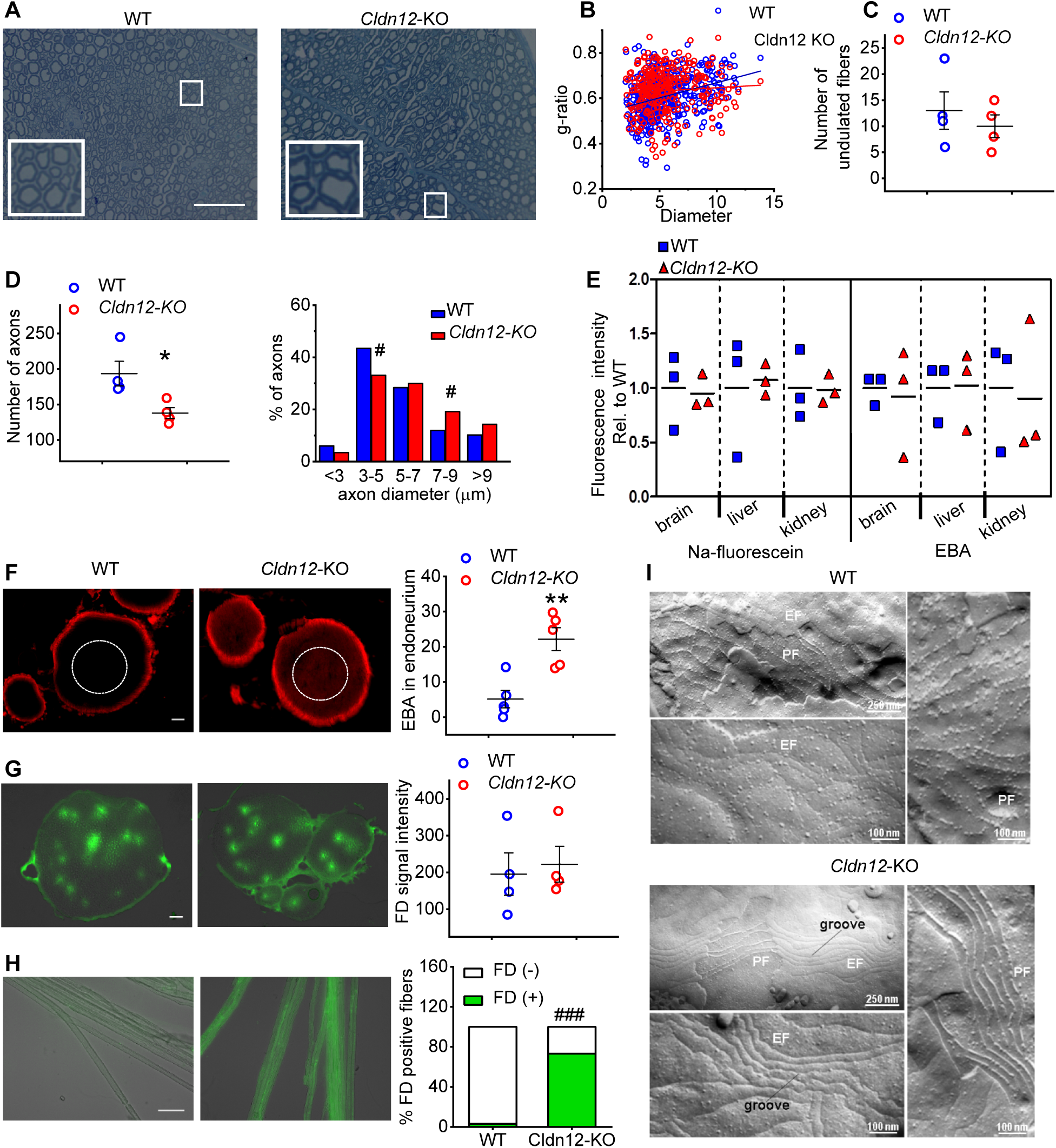
*Cldn12* deficiency results in selective nerve barrier breakdown and axonal loss. (**A**) Representative images of transverse semithin sections of the sciatic nerve from WT and *Cldn12*-KO mice including higher magnification (white square) are displayed. (**B**) The ratio of inner axonal diameter to total outer diameter of nerve fibers (g-ratio) in relation to the fiber diameter and (**C**) the number of undulated fibers. (**D**) The number of myelinated axons in the SN and distribution of the diameters of myelinated axons were quantified (*p < 0.05, one-tailed Student’s t-test; ^#^p < 0.001, Pearson’s Chi-squared test; n = 4, scale bar: 50 µm). (**E**) Uptake of Na-fluorescein (0.5 mol/kg, i.v.) and Evans blue dye associated to serum or plasma albumin (EBA, 0.26 mol/kg, i.v.; each 10 min after injection) were measured in brain, liver and kidney from WT and *Cldn12*-KO mice (two-way ANOVA, followed by Bonferroni post hoc test, n = 3). (**F-H**) Permeabilities of the blood nerve barrier (BNB) and myelin barrier in the SN were evaluated. (**F**) Examples of EBA penetration in into SN (left) and quantification (right) of fluorescence signals as measurement of the perineurial barrier of the BNB are shown (scale bar: 50 µm). (**G**) Diffusion of FD (left) and FD fluorescence was quantified in endoneurial microvessels 2 min after i.v. FD injection (right) as part of the BNB (**p < 0.01, two-tailed Student’s t-test, n (EBA) = 4, n (FD) = 5, scale bar: 50 µm). (**H**) Myelin barrier permeability was assessed by FD penetration into teased sciatic nerve fibers (left). The number of FD positive and negative fibers in the SN (right) was investigated (^###^p < 0.001, Pearson’s Chi-squared test with Yates’ continuity correction, n = 105-110 fibers from 5 mice/group, scale bar: 20 µm). (**I**) Freeze-fracture electron microscopy of the tight junction strand network was performed in the SN from WT and KO mice. Tight junction particles associated to the strands of the protoplasmic face (PF) and exoplasmic face (EF) and mesh characteristics were quantified (**Supplemental Table 7**, scale bars: 250 nm and 100 nm as indicated). n = 3 images from 4 mice/group.

To test barrier function in different organs, we used Evans blue dye associating to plasma albumin (EBA, 68 kDa), FITC dextran (FD, 70 kDa) and Na-fluorescein (368 Da). *Cldn12* deficiency did not cause significant changes in barrier permeability of the brain, liver and kidney (**Fig 4E**). EBA immersion of the SN resulted in WT in bright red fluorescence only in the EPN of WTs. *Cldn12* deficiency significantly increased the red fluorescence signal intensity within the nerve (**Fig 4F**) similar to perineurial leakage after CCI (Reinhold *et al*., 2019). No barrier breakdown was observed in endoneurial vessels after i.v. FD (**Fig 4G**). Using the FD immersion method for myelin barrier permeability (Guo *et al*., 2014), we found that the fluorescence signal significantly accumulated inside of teased nerve fibers from male *Cldn12*-KO mice demonstrating a leaky myelin barrier (**Fig 4H**). In contrast, no changes in BNB and myelin barrier permeability were noticed in fertile female *Cldn12*-KO mice (**Supplemental Fig S4B**).

In the freeze-fractured electron microscopic analysis, a significant increase in membrane-associated particle density was observed in the protoplasm-face (PF) in the SN of *Cldn12*-KO compared to WT mice (**Fig 4I**). In contrast, *Cldn12* deficiency resulted in a significant reduction in the exoplasm-face (EF) associated particle density (**Supplemental Table 7**). Quantification of tight junction structure also revealed a significant increase in mesh length and decrease in mesh diameter in the PF and EF of male *Cldn12*-KO mice. In summary, deletion of *Cldn12* causes an abnormal myelin barrier and perineurium permeability associated with alterations in tight junction morphology in the PNS with no effect on other barriers like the BBB.

### No major tight junction protein interactions nor barrier sealing properties but rather a regulatory function of claudin-12

Since tight junction proteins form strands by inter-claudin interactions (Cording *et al*., 2013), we further studied whether claudin-12 interacts with other tight junction proteins in an *in vitro* system. Tight junction-free human embryonic kidney cells (HEK)-293 cells were transfected with yellow fluorescence protein (YFP)- or turquoise (TRQ)-tight junction protein constructs (**Fig 5A-D**). Claudin-12 integrated into the plasma membrane but no homophilic *trans*-interactions were made (**Fig 5B, D**). Claudin-12 exhibited low grade heterophilic *trans*-interactions with occludin (**Fig 5C, D**). However, no heterophilic *cis*-associations of claudin-12 with PMP22 or claudin-19 were observed (**Fig 5E**). Functionally in vitro, the transepithelial resistance (TER) indicating barrier sealing remained unchanged in YFP-*Cldn 12* transfected cells compared to the positive control, YFP-*Cldn5* transfected cells (**Fig 5F**).

**Figure 5.**
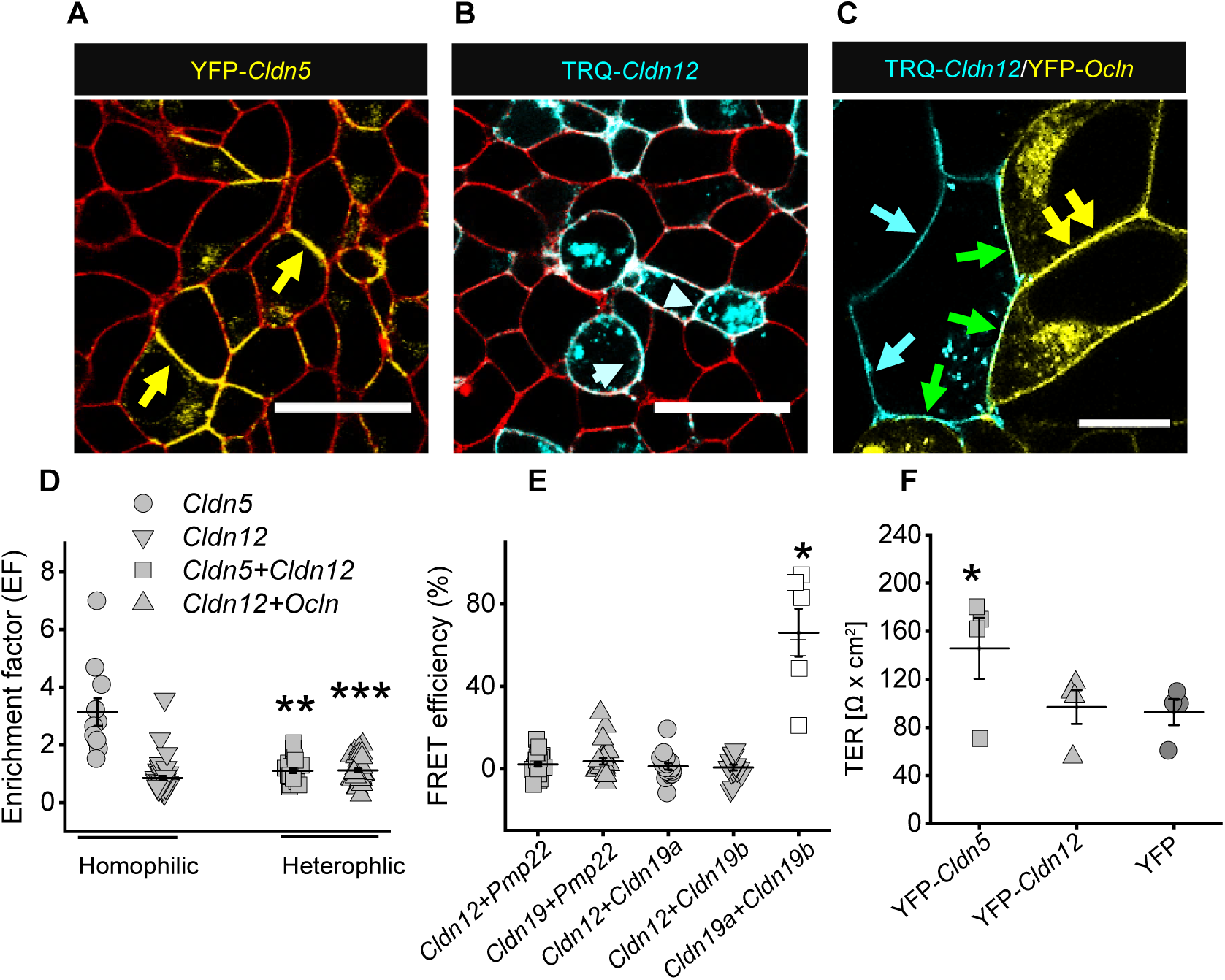
No major heterophilic or homophilic cis/trans interaction, nor an increase in transcellular electrical resistance (TER) of claudin-12. Tight junction-free HEK-293 cells transfected with yellow fluorescence protein (YFP)- or turquoise fluorescence protein (TRQ)-tagged claudin and occludin are shown, respectively. (**A**) Yellow arrows indicate claudin-5 enriched in tight junction between transfected cells (red color: trypane blue as membrane marker; live-cell imaging, scale bar: 25 µm). (**B**) TRQ-Cldn12 distribution was analyzed in cell contacts. Blue arrowheads indicate partial enrichment in tight junctions. (**C**) Representative images with double transfection are displayed. Green arrows depict contacts between claudin-12 and occludin in the tight junction. Open blue arrows show membrane location of claudin-12 in contacts with non-transfected cells. Yellow arrows indicate contacts between occludin expressing cells. (**D**) The enrichment factor (EF) between TRQ-Cldn12 in the tight junctions (EF∼1), positive control YFP-Cldn5 (EF>1, homophilic trans-interaction between neighboring membranes; n = 5) and TRQ-Cldn12/YFP-Ocln/YFP-Cldn5 heterophilic trans-interactions were quantified (*p < 0.05, compared to claudin-12, Kruskal-Wallis-test and Dunn’s multiple comparison post hoc test, n > 15). (**E**) Förster resonance energy transfer (FRET) analysis efficiencies measured cis-interaction between claudin-12 and PMP22 (n = 25), claudin-19b and PMP22 (n = 24), claudin-12 and claudin-19a (n = 17) or claudin-19b (n = 15) (single sample student’s t-test with post hoc Bonferroni-Holm analysis). FRET analysis efficiency between claudin-19a and claudin-19b is displayed as positive control (n = 6, single sample student’s t-test with post hoc Bonferroni-Holm analysis; *p < 0.05). (**F**) Measurement of TER in MDCK-II cells transfected with YFP-Cldn5 (positive control), YFP (vector control), and YFP-Cldn12 (*p < 0.05, one-tailed Student’s t-test, n = 3). Data are shown as mean ± SEM.

Since claudin-12 seemed not directly involved in barrier sealing, we next asked whether nerve injury further affects mRNA expression of known sealing tight junction proteins in the dSN or EPN of WT and *Cldn12*-KO mice. In naïve mice, *Cldn1* mRNA levels in the EPN as well as *Cldn19* mRNA levels in the dSN were significantly decreased in male *Cldn12*-KO mice (**Fig 6A-E**). *Cldn5* and *ZO-1* were unaffected. In contrast, we observed a 3.5-fold upregulation of occludin. Nerve injury caused barrier breakdown and a reduction of *Cldn1*, *Cldn5*, *Cldn19* and *ZO-1* mRNA levels in KO and WT mice by 40-80% mice. In contrast, in isolated brain capillaries no significant differences in tight junction protein mRNA expression, e.g. *Cldn1*, *Cldn3*, *Cldn5* and *Ocln* mRNA, were detected between male naïve WT and *Cldn12*-KO mice (**Fig 6F**). In line with barrier function and nociceptive threshold determination, no tight junction protein mRNA changes were observed in fertile female *Cldn12*-KO mice compared to WT mice (**Supplemental Fig. S4C, D**). Specifically, we did not observe a significant downregulation of *Cldn19*. To determine whether the proalgesic cytokine TNF-α contributes to mechanical hypersensitivity in *Cldn12*-KO mice, we examined *Tnf* mRNA expression before and after CCI in the mice. We found a strikingly high expression of *Tnf* in *Cldn12*-KO mice that was similar to the expression after CCI in the WT and KO mice (**Fig 6G**).

**Figure 6.**
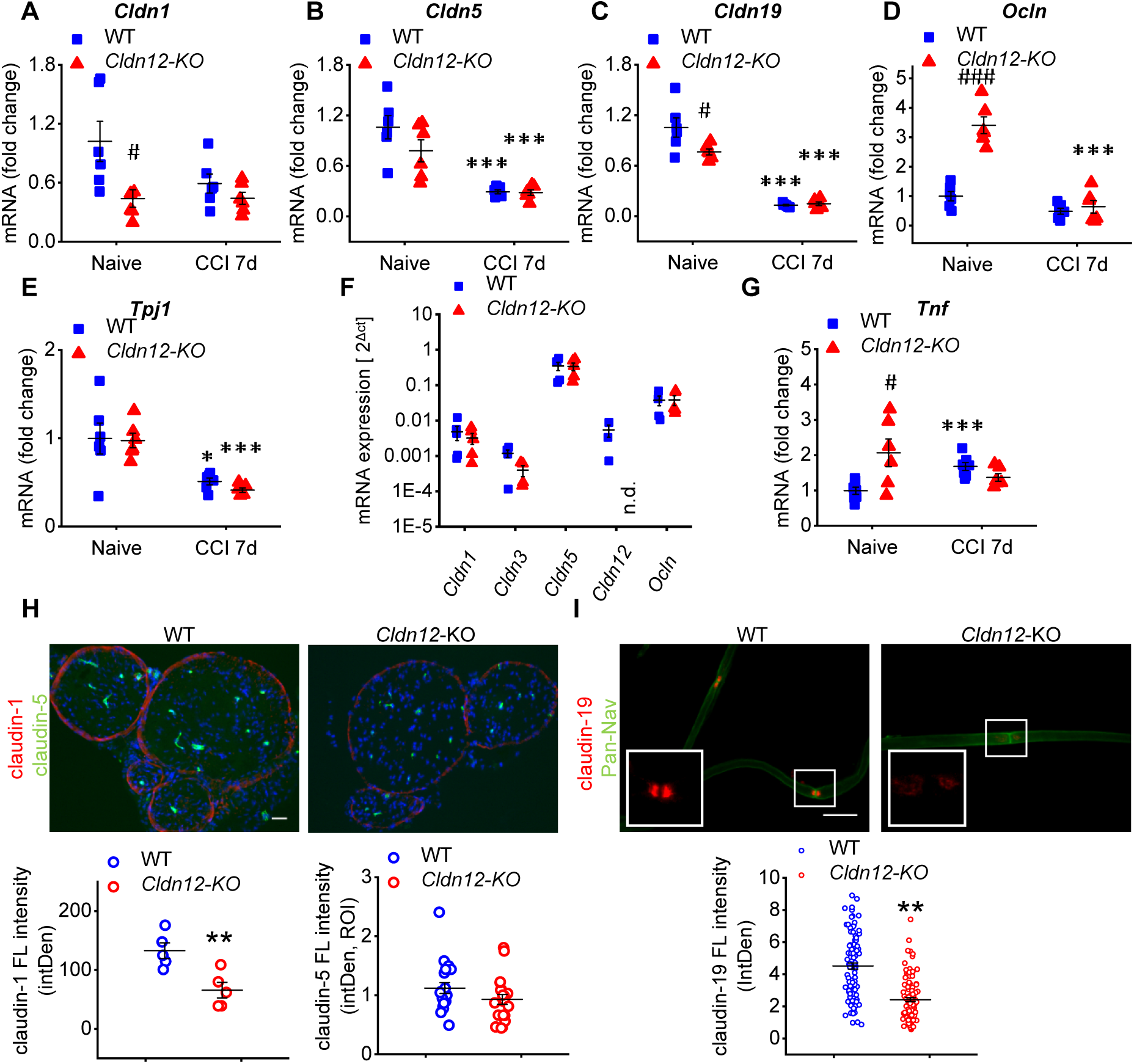
Reduction of *Cldn1* and *Cldn19* mRNA and protein in the sciatic nerve (SN) in *Cldn12*-KO mice. The mRNA expression of *Cldn1* in the epiperineurium (EPN) (**A**), *Cldn5* (**B**) *Cldn19* (**C**), *Ocln* (**D**) and *ZO-1* (*Tjp1*) (**E**) in the desheathed sciatic nerve (dSN) was quantified in WT or *Cldn12*-KO mice before and 7 d after CCI (#p < 0.05, ###p < 0.001, naïve *Cldn12*-KO versus naive WT. *p < 0.05, ***p < 0.001, CCI versus corresponding cohorts from naive WT, n = 5-6). (**F**) *Cldn1*, *Cldn3*, *Cldn5*, *Cldn12* and *Ocln* mRNA expression was analyzed in isolated brain capillaries of *Cldn12*-KO and WT mice (n = 4-5, non-detectable, n.d.). (G) *Tnf* mRNA expression in the desheathed sciatic nerve (dSN) from WT or *Cldn12*-KO mice before and 7 d after CCI. (#p < 0.05, naïve Cldn12-KO versus naive WT. ***p < 0.001, CCI versus corresponding cohorts from naive WT, n = 6-7) (**H**) Examples of immunostainings for claudin-1 (red) and claudin-5 (green) in the SN are displayed and quantified in naïve *Cldn12*-KO and WT (bottom, n = 4, scale bar: 50 µm). (**I**) Representative images of claudin-19-IR and quantification (bottom) in paranode of teased nerve fibers are shown. Nodal regions were identified with a anti pan-sodium channel Ab (pan-Nav). All above two-tailed Student’s t-test. **p < 0.01, naïve KO nerve fibers versus nerve fibers from naive WT, n = 101-104 nerve fibers from four mice per group. Data are presented as mean ± SEM. Scale bar: 20 µm.

Using immunostaining, we examined claudin-1-, claudin-5- and claudin-19-IR in the SN of male KO and WT mice. Claudin1-IR, which was mainly detected in the EPN, was significantly decreased in male *Cldn12*-KO mice (**Fig 6H**). No significant changes in claudin-5-IR were detected in endothelial cells of endoneurial vessels of KO mice (**Fig 6H**). Furthermore, *Cldn12* deficiency resulted in a significant downregulation of claudin-19-IR in the myelin barrier compared to WT mice (**Fig 6I**). As observed with mRNA levels, tight junction protein-IR were comparable in fertile female *Cldn12*-KO and WT mice (**Supplemental Fig S4C, D**). In summary, *Cldn12* deficiency in male mice attenuates mRNA/protein expression of other specific tight junction proteins sealing the perineurium and the myelin barrier, but not microvessels in the SN and the brain pointing towards a regulatory function in the PNS.

### Mechanical allodynia and myelin barrier breakdown after local suppression of *Cldn12* transcription in the SN

We next hypothesized that the mechanical allodynia from global *Cldn-12* deficiency was a consequence of claudin-12 loss specifically in the SN. To test this hypothesis, we short-term silenced *Cldn12* expression in the SN by siRNA. Local *Cldn12*-siRNA treatment downregulated *Cldn12* mRNA and was sufficient to induce myelin barrier breakdown and mechanical allodynia (**Fig 7A, B**). Similarly, local *Cldn19*-siRNA application as a positive control resulted in mechanical hypersensitivity and myelin barrier leakiness with reduced *Cldn19* mRNA in the dSN (**Fig 7A, C**). As a negative control, we applied *Clnd5*-siRNA, because claudin-5 was unaltered in *Cldn12*-KO mice. The treatment of SN with *Cldn5*-siRNA reduced *Cldn5* mRNA but did not cause myelin barrier collapse or mechanical allodynia (**Fig 7A, D**). These results demonstrate that loss of *Cldn12* or *Cldn19* in the SN was obligatory to the development of mechanical allodynia and myelin barrier leakiness.

**Figure 7.**
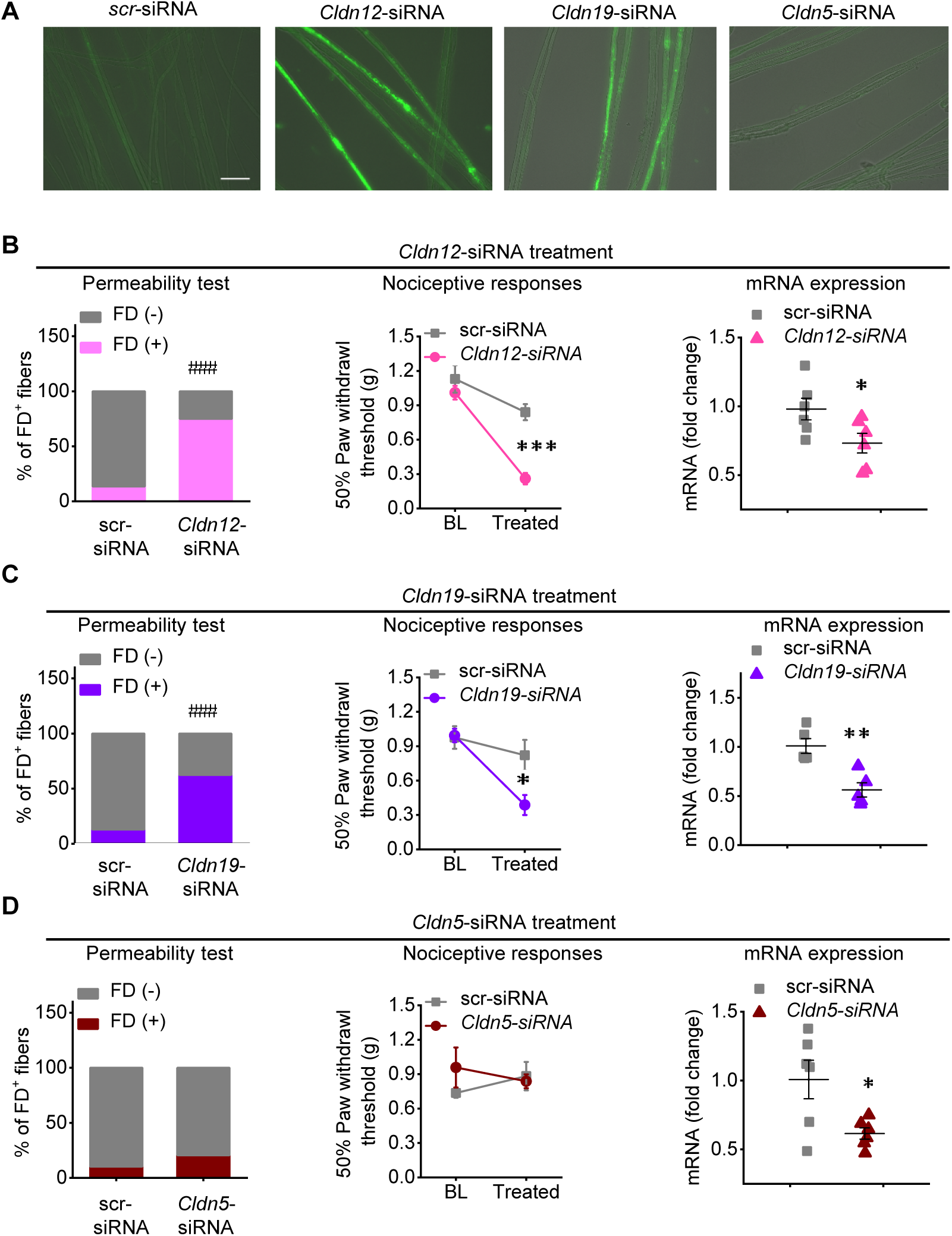
Mechanical allodynia and myelin barrier breakdown after *Cldn12* suppression. (**A-D**) Mice were treated with a single perisciatic injection of scrambled (scr), *Cldn12*-, *Cldn19*- and *Cldn5*-siRNA. (**A**) Desheathed sciatic nerves (dSN) were incubated with FICT-dextran (FD) *ex vivo* after respective *in vivo* siRNA treatment. Representative images of teased fibers are displayed (scale bar: 50 µm). Ratios of FD positive fibers from *Cldn12*- (**B**), *Cldn19*- (**C**), *Cldn5*- (**D**) or scrambled-siRNA treated sciatic nerve (left panel) were quantified. Mechanical nociceptive thresholds (middle panel, baseline, BL, and after siRNA application) and mRNA expression of *Cldn12*, *Cldn19* and *Cldn5* in the SN (right panels) were analyzed following the siRNA injection. (*p < 0.05, **p < 0.01, ***p < 0.001, *Cldn*-siRNA versus scrambled-siRNA, two-tailed Student’s t-test, n = 5-6; ###p < 0.001, Pearson’s Chi-squared test with Yates’ continuity correction, n = 78-118 fibers from 4-5 mice per groups). Data are presented as mean ± SEM.

### Impaired expression of SHH, a master barrier stabilizer, in male *Cldn12*-KO mice

Because Schwann cells forming the myelin barrier were affected in *Cldn12*-KO, we further explored myelin associated proteins and transcription factors important for Schwann cell differentiation. The expression of Myelin Basic Protein (*Mbp*) mRNA, a structural protein in myelinating Schwann cells, was unaltered in the dSN of *Cldn12*-KO mice (**Fig 8A**) although these mice had a reduced number of axons (**Fig 4C**). In contrast, *Pmp22* mRNA was downregulated in *Cldn12*-KO (**Fig 8B**). PMP22 is a transmembrane protein related to tight junction proteins, necessary for compact myelin formation and altered in certain forms of hereditary neuropathy (Guo *et al*., 2014) The Schwann cell transcription factors HMG-box 10 (*Sox10*) and growth response protein 2 (*Krox20*) play a role in nerve injury and are necessary for nerve regeneration after nerve injury (Quintes and Brinkmann, 2017). No differences of *Sox10* and *Krox20* mRNA were observed between the genotypes indicating no general defect in Schwann cell differentiation (**Fig 8C, D**).

**Figure 8.**
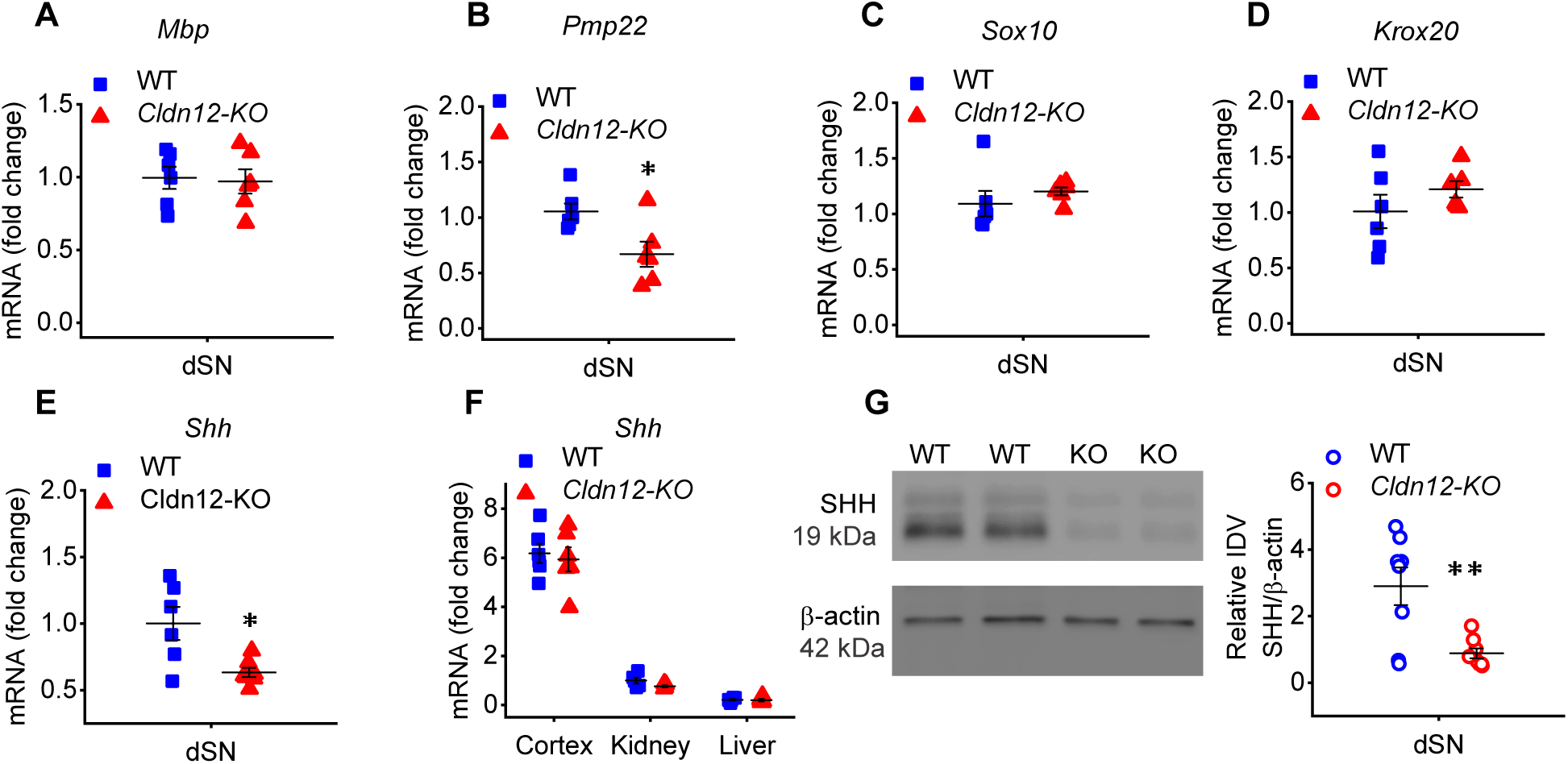
*Cldn12* deficiency results in downregulation of *Shh* expression in the SN, but not in the brain cortex, liver and kidney. (**A**) The mRNA expression of myelin basic protein (*Mbp*), (**B**) peripheral myelin protein 22 (*Pmp22*) and the two transcription factors (**C**) *Sox10*, (**D**) *Krox20* and (**E**) *Shh* were measured in the dSN from WT and *Cldn12*-KO mice. The mRNA expression was always expressed as fold change to naïve WT mice. (**F**) The *Shh* mRNA was quantified in the brain cortex, kidney and liver from WT and KO mice. (**G**) Representative images of Western blot analysis of SHH protein expression in the dSN from WT and KO mice. Bands were analyzed by densitometry (intensity density value, IDV). Uncut blots are found in **Supplemental Fig S5**. *p < 0.05, ** p < 0.01, naïve *Cldn12*-KO versus naïve WT, all two-tailed Student’s t-test, PCR n = 6, Western blot n = 8). Data are presented as mean ± SEM.

Neuronal barriers are stabilized by the morphogen SHH (Alvarez *et al*., 2011). In the dSN from *Cldn12*-KO mice, *Shh* mRNA and SHH protein were significantly downregulated in naïve animals (**Fig 8E, G, H**). In contrast, *Shh* mRNA expression was normal in the cortex, liver and spleen of KO mice in accordance with unaltered tight junction protein mRNA expression in these tissues (**Fig 8F**). In the same line, *Shh* mRNA in fertile female mice was unaffected (**Supplemental Fig S4E, F**). Collectively, these results demonstrate that *Cldn12* deficiency is associated with a reduction of SHH causing tight junction protein loss in the SN and myelin barrier only in the PNS, which lowers mechanical nociceptive thresholds in male animals (**Fig. 9**).

**Figure 9.**
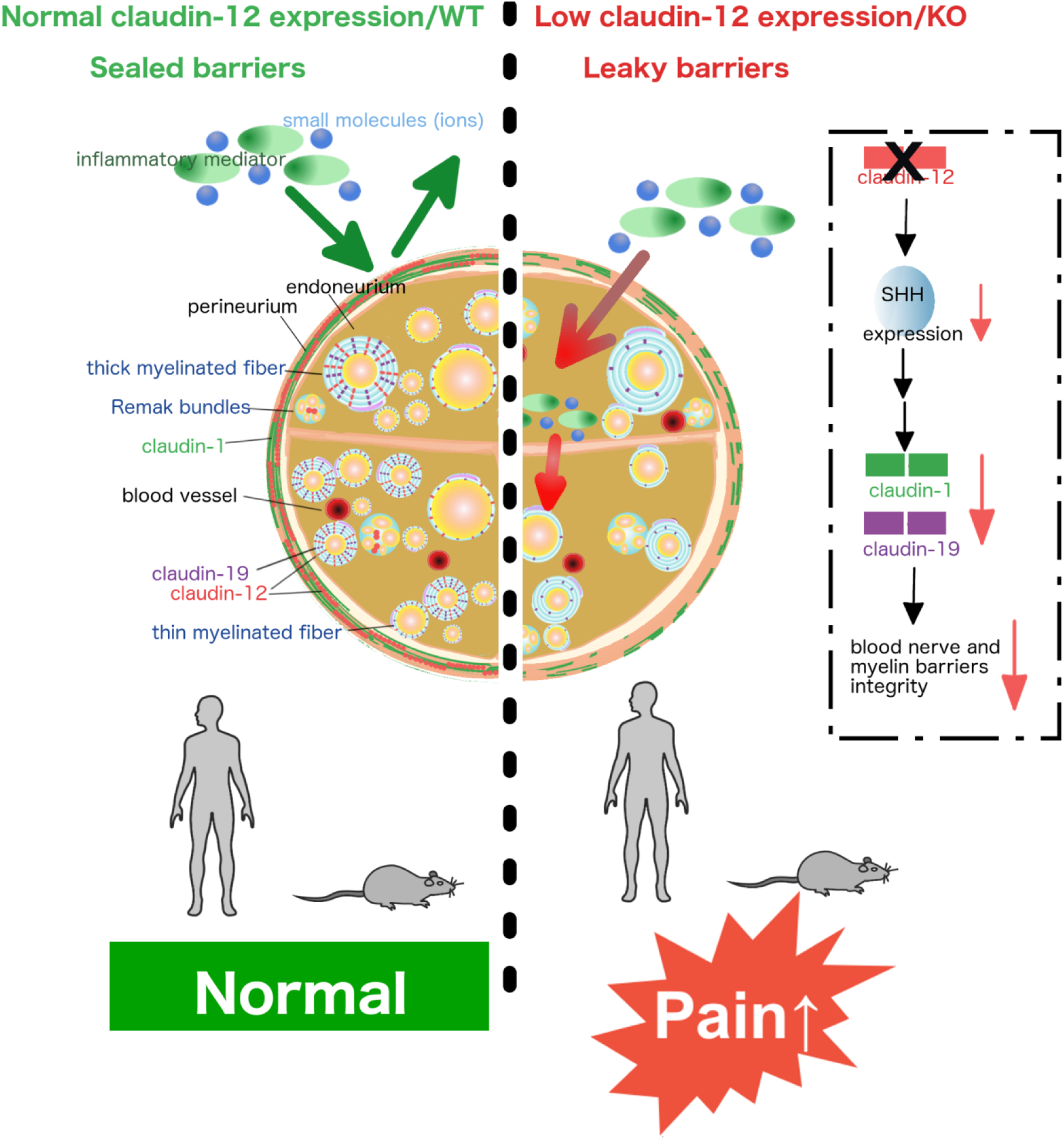
Role of claudin-12 on nerve barriers sealing and pain. Drawing depicts the possible cellular and molecular events in the presence (left) and absence of claudin-12 (right). Reduced claudin-12 or KO increases the permeability of the perineurial and the myelin barrier in male mice. Claudin-12 KO is accompanied by reduced levels of the morphogen SHH thereby destabilizing barriers via reduced expression tight junction protein networks including claudin-1 and claudin-19.

## Discussion

In this study, we investigated the newly-discovered tight junction protein claudin-12 in peripheral nerves of patients, animals and in transfected cells. Claudin-12 was significantly lost only in male and female postmenopausal patients with painful CIDP or non-inflammatory PNP compared to the non-painful group. Male naïve and nerve-injured *Cldn12*-KO mice were hypersensitive to mechanical nociceptive stimuli and suffered from a breakdown of the perineurial barrier and myelin barrier in the PNS (**Fig. 9**). *Cldn12* deficiency lowered mRNA and protein expression of other tight junction proteins namely claudin-1 and -19, and the number of myelinated axons. Loss of claudin-12 in the SN also reduced the morphogen and barrier stabilizer SHH explaining the effect on other tight junction proteins. The regulatory and barrier stabilizing role of claudin-12 was confirmed by local siRNA treatment. Together, our results suggest, for the first time, that *Cldn12* expression in the PNS regulates barrier permeability and mechanical nociception via other tight junction proteins such as claudin-19, claudin-1 and SHH in male but not fertile female mice.

Low amounts of cldn-12 are expressed in microvascular cells in the brain (Ohtsuki *et al*., 2007; Castro Dias *et al*., 2019b), lung (Chen *et al*., 2018), intestine (Chen *et al*., 2018), SN (Shimizu *et al*., 2008) and cancer cells (Yang *et al*., 2015). Besides controlling Ca^2+^ absorption in the gastrointestinal system, the function of claudin-12 in the nervous system is unknown. It was previously postulated that claudin-12 is important for BBB formation. Since all neurological tests except for mechanical nociceptive threshold were normal in naïve *Cldn12*-KO mice in our study, *Cldn12* seems to imply at least not to be crucial for CNS function under normal conditions. This phenotype was replicated in a recent study (Castro Dias *et al*., 2019a) after our study was in preprint.

Tight junction proteins are differentially expressed throughout peripheral nerves (Yang *et al*., 2016; Reinhold and Rittner, 2017). For example, claudin-1 is the major protein in the perineurium (Hackel *et al*., 2012; Sauer *et al*., 2014). Claudin-5 is mainly expressed in the endothelial layer of blood vessels in the SN regulating this part of the BNB (Moreau *et al*., 2016; Moreau *et al*., 2017a). Claudin19 is found in the paranode of nerve fibers (Miyamoto *et al*., 2005; Guo *et al*., 2014). Claudin-12-IR as well as *Cldn12* mRNA were found in myelinating and non-myelinating Schwann cells and the perineurium. Specificity of the claudin-12 IR in Schwann cells was confirmed repeatedly in *Cldn12*-KO in contrast to the murine skeletal muscle in other studies (Castro Dias *et al*., 2019a). *Cldn12* deficiency suppressed *Cldn1* in the EPN and *Pmp22* and *Cldn19* expression in the dSN in male mice. In line with this, perineurium and myelin barrier were more permeable to medium-sized dyes. *Cldn5* in endoneurial vessels and its barrier function were unaltered. *Ocln* mRNA was upregulated, but this seemed to have no functional consequence. Thus, the *Cldn12* deficiency-induced leaky barrier in the SN may derive from the downregulation of claudin-1 in the perineurium and claudin-19/PMP22 in the dSN, suggesting that claudin-12 could be involved in the integrity of perineurial barrier of the BNB and myelin barrier. The presented data also imply that *Cldn12* has a more prominent role in the PNS, since we observed no difference in mRNA expression of *Cldn1*, *Cldn5* and *Cldn19*, or barrier function in the brain, liver and kidney of male *Cldn12*-KO mice. In addition, tight junction ultrastructure in the perineurium of claudin-12-free peripheral nerves was altered. This finding would explain increased tracer uptake into nerve fibers, and that the deficiency of *Cldn12* causes loss of tightening and particle forming tight junction proteins, as for instance of claudin-1 (Cording *et al*., 2013) or claudin-19 (Miyamoto *et al*., 2005). Therefore, a general regulatory role of claudin-12 is assumed in peripheral nerves.

The morphogen SHH is known to regulate barrier tightness in the BBB and the BNB. In the BNB, *Shh* is upregulated after nerve injury as a compensatory mechanism (Moreau *et al*., 2016). Repressing the hedgehog/smoothened (smo) pathway via the smo receptor antagonist cyclopamine suppresses *Cldn5* and *Ocln* mRNA expression in the SN and elicits mechanical hypersensitivity (Moreau *et al*., 2016; Moreau *et al*., 2017a; Moreau *et al*., 2017b). We did not see downregulation of *Cldn5* and *Ocln* mRNA in male *Cldn12*-KO mice but rather *Cldn19*, *Cldn1* and *Pmp22* mRNA. At least *Pmp22* is also induced by SHH *in vitro* (Ingram *et al*., 2002). It is therefore conceivable that SHH regulates other tight junction proteins in peripheral nerves as well. In our study, *Cldn12* deletion reduced *Shh* expression exclusively in the SN in male but not in fertile female mice. Also, *Shh* expression in the brain, liver and kidney of *Cldn12*-KO mice were unaffected. Interestingly, *Shh* expression can be stimulated by estrogen in estrogen receptors positive gastric cancer cells (Kameda *et al*., 2010). After nerve injury, estradiol improves nerve recovery, at least in part, by increasing SHH signaling and SHH-induced angiogenesis after crush injury. This is not a direct effect but rather due to a reduced expression of the SHH inhibitor hedgehog-interacting protein (HIP) in endothelial and Schwann cells (Sekiguchi *et al*., 2012). Thus, our data point at a role for claudin-12 in setting neuronal barrier function and imply that in its absence and with low estrogen SHH is downregulated selectively in the PNS in mice. This ultimately reduces the expression of selected tight junction proteins in the peripheral nerve forming the BNB and myelin barrier.

Deletion of *Cldn12* gene via KO or siRNA in the SN led to mechanical allodynia. Barrier breakdown could result in the dysregulated entrance of ions as well as pro-nociceptive and inflammatory mediators (e.g. IL-1β and TNF-α) (Ji *et al*., 2016). Indeed, *Tnf* mRNA was upregulated in naïve KO mice. These could, for example, activate transient receptor potential vanilloid 1 (TRPV1) or transient receptor potential ankyrin 1 (TRPA1) expressed along the axonal membrane and thereby increase excitability causing hypersensitivity (Sauer and Reeh, 2009; Weller *et al*., 2011).

A few tight junction protein KO models collectively implicate critical roles for tight junction proteins in nociception in the PNS. For example, loss of junctional adhesion molecule-C (JAM-C, *Jam3*) in Schwann cells results in mechanical hypersensitivity (Colom *et al*., 2012). Tight junction protein expression in the SN is dependent on LRP1 (Hackel *et al*., 2012). Mice with Schwann cell-specific KO of low-density lipoprotein receptor-related protein 1 display abnormalities in axon myelination and ensheathment of axons by non-myelinating Schwann cells in Remak bundles and mechanical allodynia (Orita *et al*., 2013). Knockdown of *Cldn11* and *Cldn19* results in delayed nerve conduction velocity, abnormal behavioral responses and motor function deficiencies but nociception was not examined (Gow *et al*., 1999; Miyamoto *et al*., 2005; Maheras *et al*., 2018). Here, we now provide genetic evidence via full KO or siRNA that *Cldn12* similar to *Cldn19* are critical for the myelin barrier and normal mechanonociception.

Three tight junction proteins (claudin-1, -19 and ZO-1) were lowered in patients with CIDP or non-inflammatory PNP with high fiber loss, both in males and in postmenopausal females. This pattern of tight junction protein loss could simply be an associated sign of severe neuropathy and nerve destruction. Alternatively, early barrier breakdown might fuel the autoimmune attack in CIDP or the diffusion of other exogenous toxic mediators e.g. autoantibodies resulting in accelerated nerve destruction. However, the two groups did not differ in functional impairment measured by overall disability sum score (ODSS) arguing against a significant impact of clinical severity, at least. Kanda et al. described downregulated claudin-5 and ZO-1 in the biopsies of ten CIDP patients (Kanda *et al*., 2004). In our present larger study, tight junction protein alterations were independent of the presence of inflammation (CIDP vs. non-inflammatory PNP) and no difference in claudin-5-IR was observed. Most importantly, claudin-12 expression in Schwann cells discriminated painful from non-painful PNP. This indicates a central role of the myelin barrier in painful CIDP or non-inflammatory PNP in male and postmenopausal female patients. Future studies could specifically study premenopausal PNP and CIDP patients to test whether female sex hormones also protect patients against painfulness. However, these patients will be more difficult to recruit due to the epidemiology of PNPs.

The presented study has limitations including a small number of patients, only postmenopausal female patients, different etiologies of patients of non-inflammatory PNP and no healthy controls. Finally, barrier function using our functional assays would have been helpful to determine the significance of our findings.

In summary, loss of *Cldn12* sex-dependently causes mechanical allodynia, operating via myelin dysfunction/degradation and SHH suppression-induced loss of selected tight junction proteins in the PNS in mice. Further, claudin-12 is decreased in CIDP or non-inflammatory PNP patients with pain. Future studies will be aimed at the exact cellular function of claudin-12 in Schwann cells to understand its contribution to pain, sensory processing and barrier function. In this line, results of this study might explain the frequent side effect of pain in commonly used hedgehog/smo inhibitors for cancer treatment like sonidegib or vismodegib. Finally, pushing myelin barrier sealing via the hedgehog pathway including possible small molecules that could be used therapeutically [e.g. activation of smo with smoothened agonist (SAG) (Guo *et al*., 2014), purmorphamine (Chechneva *et al*., 2014) or oxysterols] could open new avenues in developing drug treatments for neuropathy.

## Acknowledgement

We are grateful to our patients who consented to be a part of this study. We also acknowledge the invaluable assistance in animal husbandry of the ZEMM.

We thank Jianghui Hou, Washington University, St. Louis, USA, and Nina Himmerkus, Institute of Physiology, Christian-Albrechts-University of Kiel, Germany, for providing the claudin-19 Ab. We acknowledge Roland Naumann, TCF, MPI-CBG Dresden for stem cell microinjection/transfer and Mohammed Selloum, Mouse Clinical Institute, Illkirch/France for providing us with the phenotyping data of general sensory-motor function functions of *Cldn12*-KO and WT littermates.

## Author contributions

JTC and HLR were responsible for the study concept and design. JTC, XH and IUO conducted the PNS barrier analyses and pain behavior experiments. IEB, RB, OBK generated and analyzed the claudin-12-KO. DG, LW and SD performed the in vitro experiments. PFB accomplished the tight junction morphology experiments. CS and KD conducted the clinical study. JS, AKR and XH analyzed the patients’ samples. CS, AB and IEB provided critical revision and important intellectual content. HLR, AKR and XH obtained funding for the study. JTC, MKH and HLR drafted and edited the article.

## The paper explained

### Problem

Tight junction proteins seal the blood-nerve and blood myelin barrier. However, the role the tight junction proteins claudin-12 expressed in peripheral nerves is inexplicable and lacks clinical evidence and basic scientific mechanisms.

### Results

We serendipitously found that in male or female postmenopausal patients with chronic inflammatory demyelinating polyneuropathy and non-inflammatory polyneuropathy, pain was specifically associated with reduced claudin-12 immunoreactivity in Schwann cells of the sural nerve. This led us to hypothesis that neuropathic pain arises from peripheral myelin barrier integrity breakdown due to the absence of claudin-12. We next created a germline knockout mouse line demonstrating that global deficiency in claudin-12 leads to mechanical hypersensitivity, perineurial and myelin barrier breakdown only in male mice. Furthermore, selected other tight junction proteins and the barrier-stabilizing molecule sonic hedgehog in the PNS were downregulated. Other barriers including the blood brain barrier and female mice were unaffected, suggesting a role of claudin-12 in maintaining peripheral myelin barrier integrity.

### Impact

Our work indicates that claudin-12 is a regulatory tight junction protein for the myelin barrier in Schwann cells controlling mechanical nociceptive pain sensation sex-dependently in patients and mice. This study further implies tha claudin-12 and the SHH pathway could be an attractive target for novel neuropathic pain therapies protecting myelin barriers.

---

**Supplemental Figure 1.**
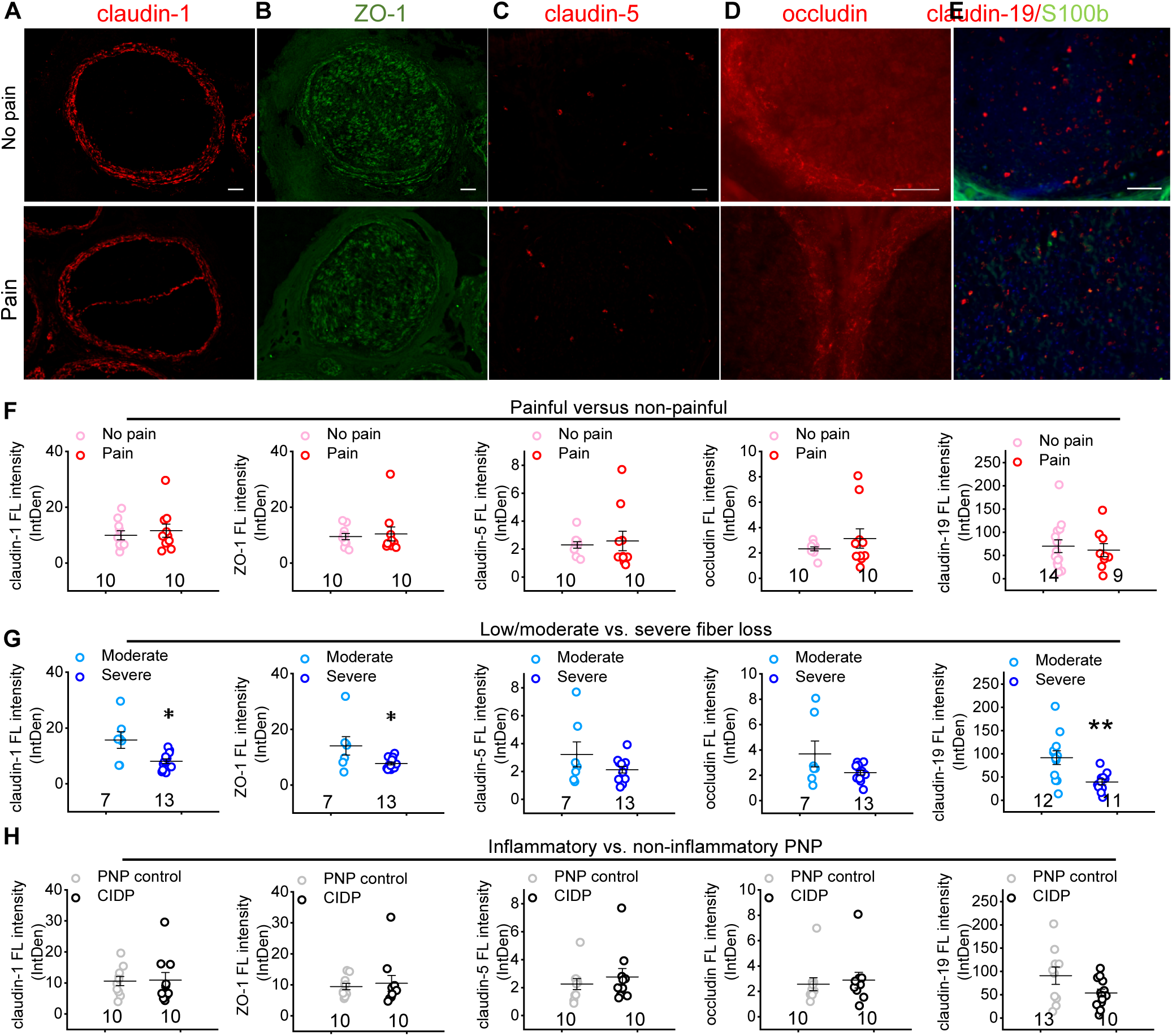

**Supplemental Figure 2.**
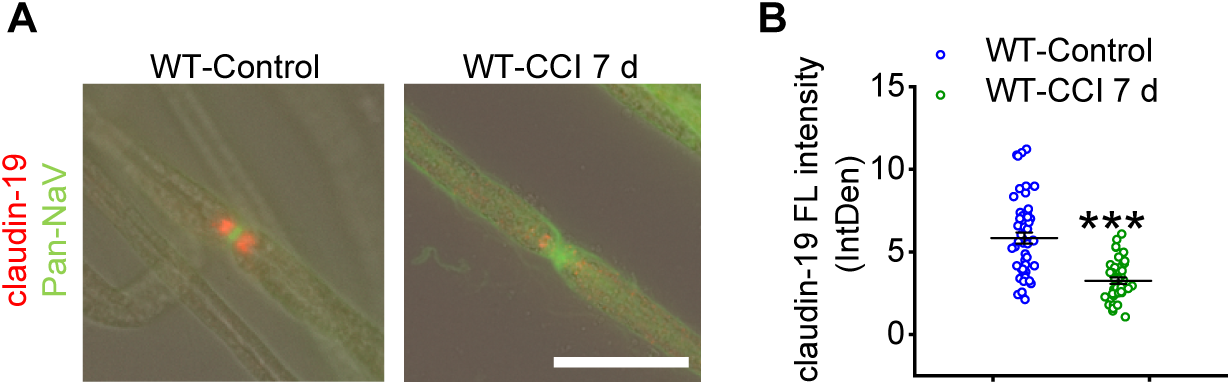

**Supplemental Figure 3.**
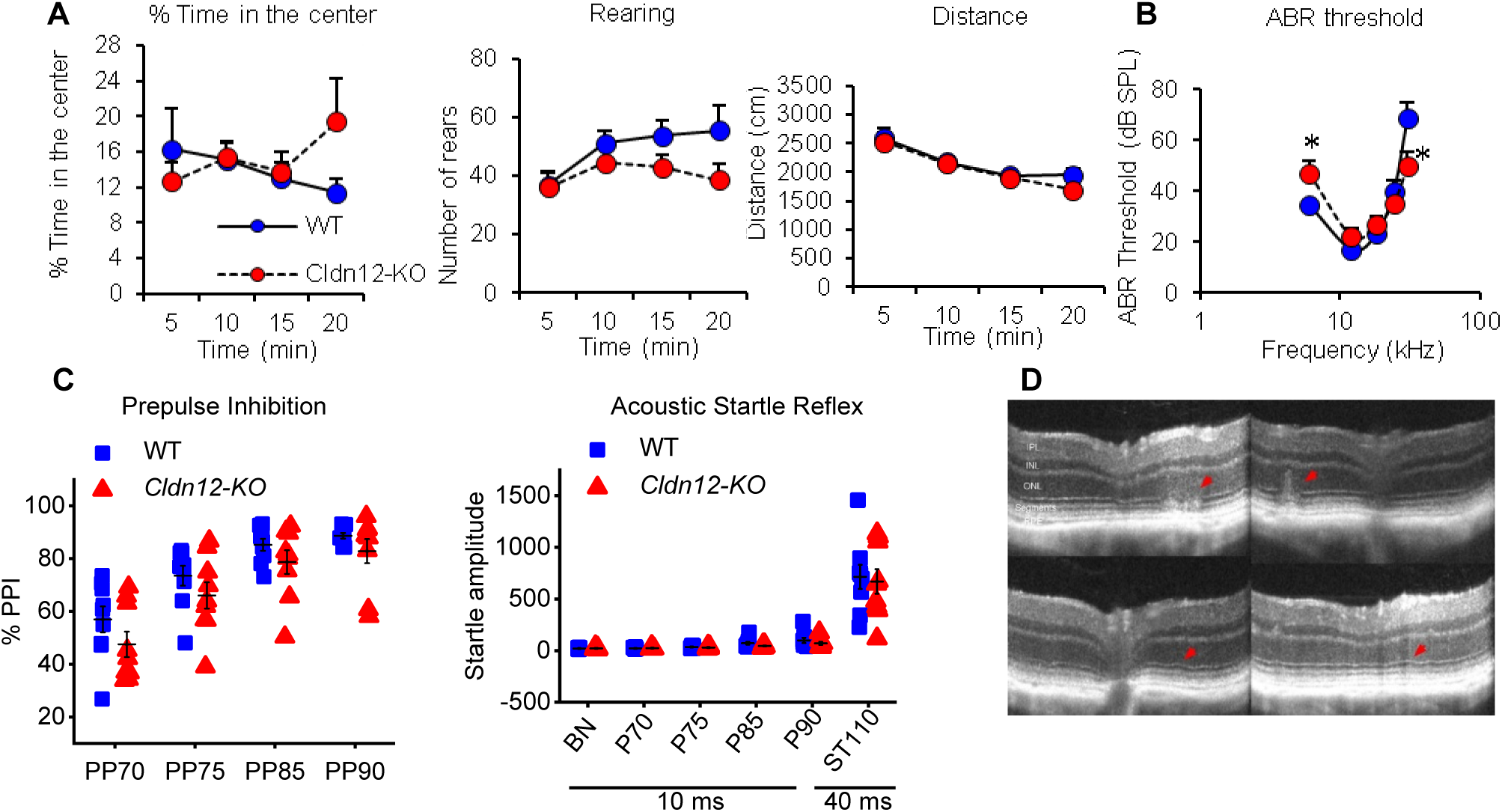

**Supplemental Figure 4.**
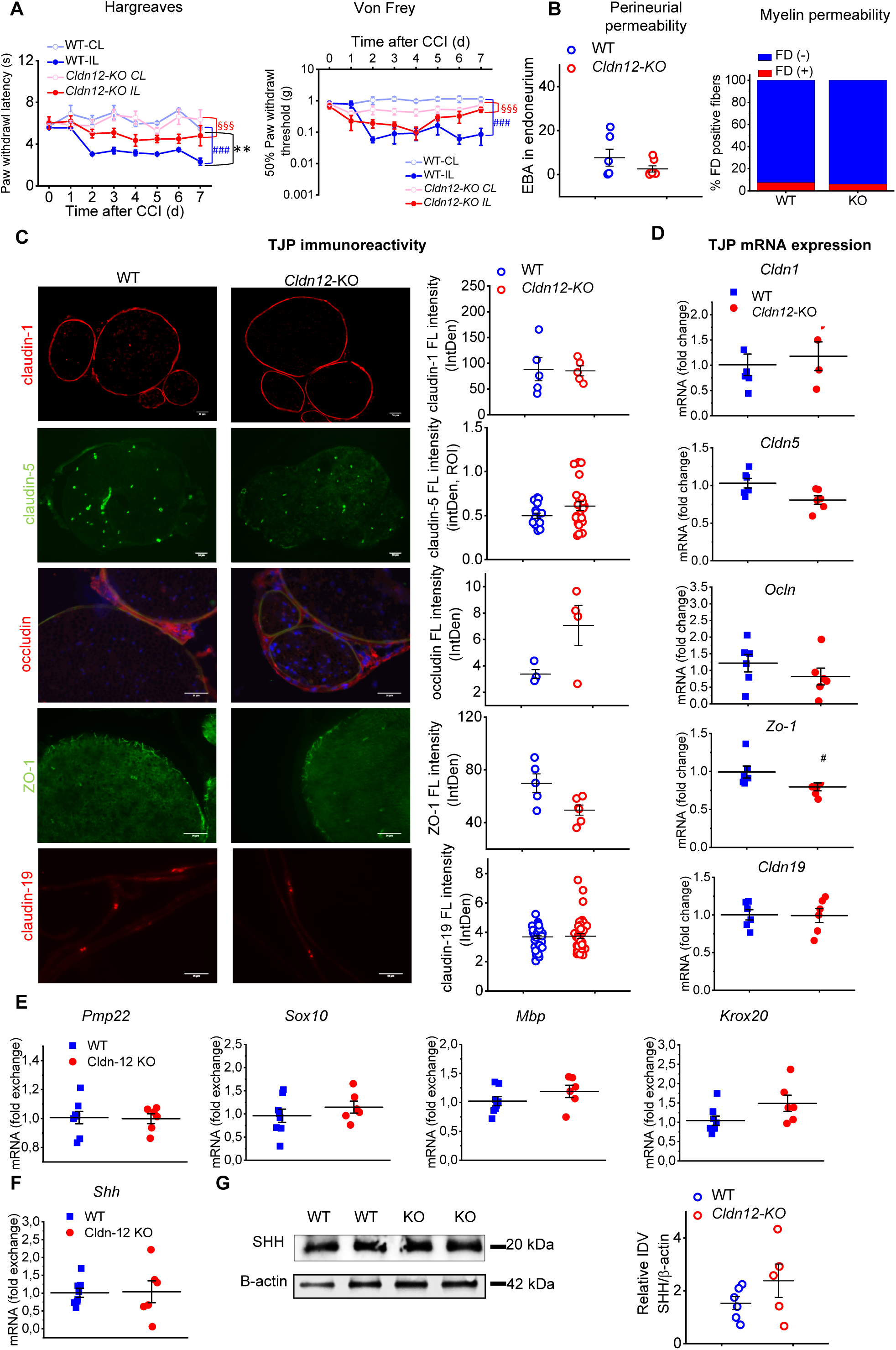

**Supplemental Figure 5.**
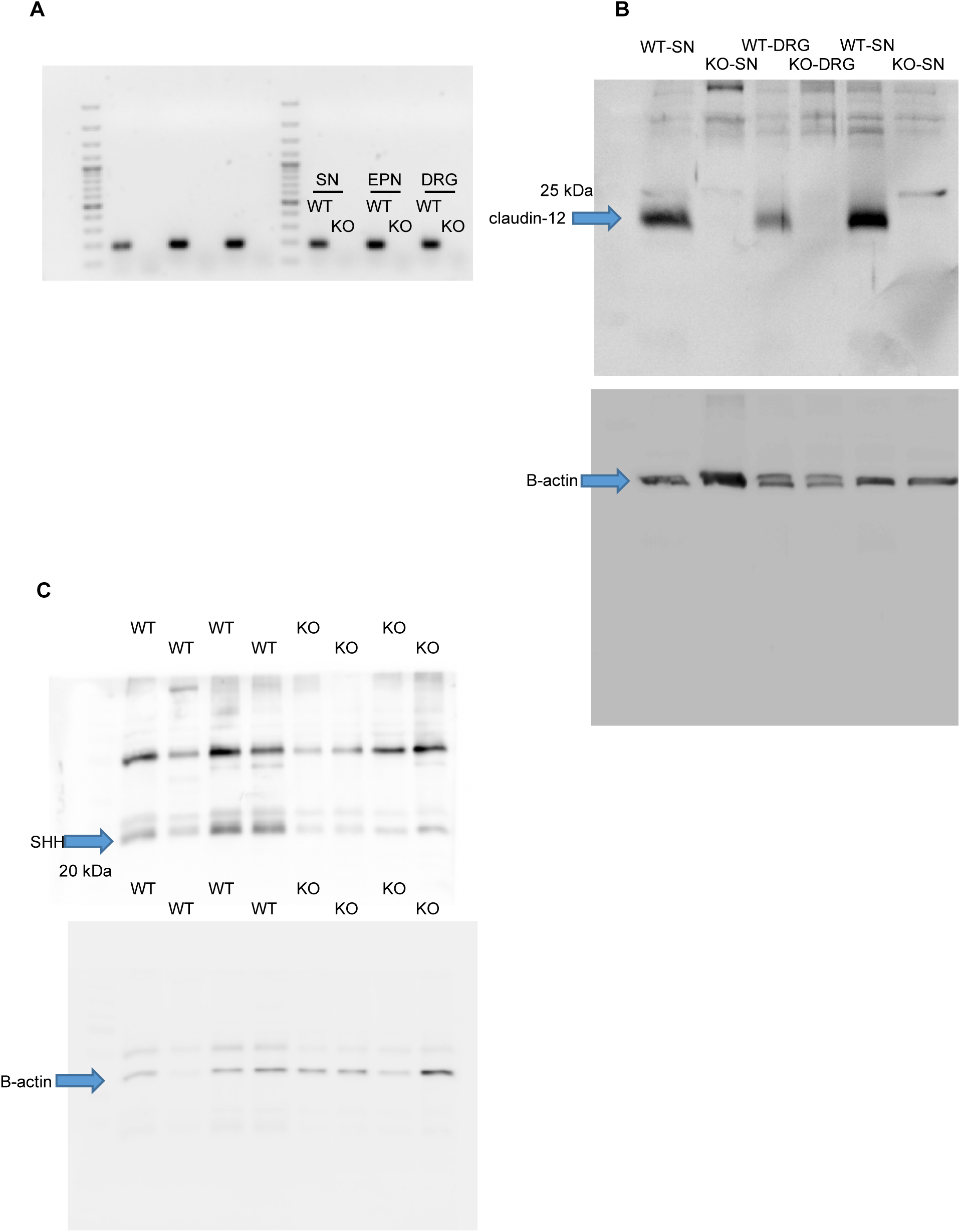
Entire unedited gel (A) Full unedited gel for Figure 2F (B) Full unedited gel for Figure 2G. (C) Full unedited gel for Figure 8G.

