## supplemental tables and figures for "Claudin-12 deficiency causes nerve barrier breakdown, mechanical hypersensitivity and painfulness in polyneuropathy"

**Chen et al.**

**Supplemantary information**

- **Supplemental Table 1.** Clinical features of chronic inflammatory demyelinating polyneuropathy (CIDP) and non-inflammatory polyneuropathy (PNP) (Supplemental Figure 1A-D)

#### - Supplemental Table 2. Clinical features of PNP patients divided into low to moderate or severe fiber loss (Supplemental Figure 1E)

#### - Supplemental Table 3. Clinical features of PNP patients divided into pain and no pain (Figure 1)

#### - Supplemental Table 4. List of RNAscope probes used in in-situ hybridization

#### - Supplemental Table 5. List of primers and Taqman probes used in qRT-PCR analysis

#### - Supplemental Table 6. List of primary antibodies used in IHC and Western blotting

#### - Supplemental Table 7. Quantitation of the tight junction network (Figure 4F)

#### - Supplemental Methods

- **Supplemental references**

**- Supplemental Figure S1.** Reduced immunoreactivity of claudin-1, -19 and ZO-1 but not claudin-5 or occludin in patients.

- **Supplemental Figure S2.** Loss of claudin-19 in the paranode after chronic constriction injury (CCI).

- **Supplemental Figure S3.** No differences in anxiety-related behavior, sensorimotor gating and auditory function of *Cldn12*-KO mice.

- **Supplemental Figure S4.** No significant difference in thermal and mechanical nociceptive responses and barrier permeability are observed in female Cldn12-KO mice.

#### Supplemental Table 1. Clinical features of chronic inflammatory demyelinating polyneuropathy (CIDP) and non-inflammatory polyneuropathy (PNP) (Supplemental Figure 1A-D)

| **Characteristic** | **CIDP**  **(n = 10)** | **non-inflammatory PNP**  **(n = 10)** | **p-value** |
| --- | --- | --- | --- |
| **Age** (years) | 68 ± 2 | 68 ± 2 | p > 0.05 |
| **Sex ratio** (M/F) | 7/3 | 5/5 | p > 0.05 |
| **Duration of disease** (months) | 37 ± 12 | 60 ± 14 | p > 0.05 |
| **ODSS** | 3 ± 0.6 | 1.9 ± 0.2 | p > 0.05 |
| **Fiber loss** |  |  | p > 0.05 |
| **moderate/low** | 2 | 5 |  |
| **severe** | 8 | 5 |  |
| **Pain** |  |  | p > 0.05 |
| **yes** | 6 | 5 |  |
| **no** | 4 | 5 |  |
| **Histology** |  |  | p > 0.05 |
| **axonal** | 3 | 6 |  |
| **demyelinating** | 7 | 3 |  |
| **mixed** | 0 | 1 |  |

All data are shown as mean ± SEM. Two-tailed student’s t-test and Chi-Square test were used to determine the statistical significance.

#### Supplemental Table 2. Clinical features of PNP patients divided into low to moderate or severe fiber loss (Supplemental Figure 1E)

| **Characteristic** | **Low and moderate fiber loss (n = 12)** | **Severe fiber**  **loss (n = 11)** | **p-value** |
| --- | --- | --- | --- |
| **Age** (years) | 68 ± 3 | 66 ± 3 | p > 0.05 |
| **Sex ratio** (M/F) | 10/2 | 7/4 | p > 0.05 |
| **Duration of disease** (months) | 42 ± 12 | 55 ± 16 | p > 0.05 |
| **ODSS** | 3 ± 0 | 3 ± 0 | p > 0.05 |
| **Diagnosis** |  |  | p < 0.01 |
| **CIDP** | 3 | 9 |  |
| **non-inflammatory PNP** | 9 | 2 |  |
| **Pain** |  |  | p > 0.05 |
| **yes** | 5 | 4 |  |
| **no** | 7 | 7 |  |
| **Histology** |  |  | p > 0.05 |
| **axonal** | 6 | 3 |  |
| **demyelinating** | 6 | 7 |  |
| **mixed** | 0 | 1 |  |

All data are shown as mean ± SEM. Two-tailed student’s t-test and Chi-Square test were used to determine the statistical significance.

#### Supplemental Table 3. Clinical features of PNP patients divided into pain and no pain (Figure 1)

| **Characteristic** | **Pain**  **(n = 12)** | **No-pain**  **(n = 11)** | **p-value** |
| --- | --- | --- | --- |
| **Age** (years) | 68 ± 2 | 67 ± 3 | p > 0.05 |
| **Sex ratio** (M/F) | 6/6 | 11/0 | p < 0.01 |
| **Duration of disease** (months) | 33 ± 6 | 60 ± 16 | p > 0.05 |
| **ODSS** | 3 ± 1 | 3 ± 0 | p > 0.05 |
| **Diagnosis** |  |  | p > 0.05 |
| **CIDP** | 6 | 6 |  |
| **non-inflammatory PNP** | 6 | 5 |  |
| **Fiber loss** |  |  | p > 0.05 |
| **severe** | 6 | 5 |  |
| **moderate/low** | 6 | 6 |  |
| **Histology** |  |  | p > 0.05 |
| **axonal** | 5 | 4 |  |
| **demyelinating** | 6 | 7 |  |
| **mixed** | 1 | 0 |  |

All data are shown as mean ± SEM. Two-tailed student’s t-test and Chi-Square test were used to determine the statistical significance.

#### Supplemental Table 4. List of RNAscope probes used in in-situ hybridization

| Gene | Cat. No. | Reference sequence |
| --- | --- | --- |
| RNAscope® mm-*Cldn12* | 520441 | NM_022890.2 |
| RNAscope® positive control | 313911 | NM_011149.2 |
| RNAscope® negative control | 310043 | EF191515 |

#### Supplemental Table 5. List of primers and Taqman probes used in qRT-PCR analysis

| Gene | Forward primers | Reverse primers |
| --- | --- | --- |
| 18S^1^ | 5’-CTCAACACGGGAAACCTCAC-3’ | 5’-CGCTCCACCAACTAAGAACG-3’ |
| Β-actin (*Actb*) | 5’- TTTGAGACCTTCAACACCCC -3’ | 5’- ATAGCTCTTCTCCAGGGAGG -3’ |
| I *Cldn1* | 5’-CTGTGGATGTCCTGCGTTTC-3’ | 5’-TTACCATCAAGGCTCGGGTT-3’ |
| II *Cldn1* | 5’-GATGTGGATGGCTGTCATTG-3’ | 5’-CGTGGTGTTGGGTAAGAGGT -3’ |
| *Cldn3* | 5’-GAGATGGGAGCTGGGTTGTA -3’ | 5’-GTAGTCCTTGCGGTCGTAGG -3’ |
| I *Cldn5* | 5’-CAGTTAAGGCACGGGTAGCA-3’ | 5’-GGCACCGTCGGATCATAGAA-3’ |
| II *Cldn5* | 5’-CTGGACCACAACATCGTGAC -3’ | 5’-GCCGGTCCAGGTAACAAAGA -3’ |
| *Cldn12* | 5’-AACTGGCCAAGTGTCTGGTC-3’ | 5’-AGACCCCCTGAGCTAGCAAT-3’ |
| *Cldn19* | 5’-GTGGATGTCTTGCGCCTCT-3’ | 5’-CTCGTGCTGACTGGATATGAC-3’ |
| *ZO-1 (Tjp1)* | 5’-TGACTCCTGACGGTTGGTCT-3’ | 5’-AGGACAGAAACACAGTTGGCT-3’ |
| I *Ocln* | 5’-GTGAGCACCTTGGGATTCCG-3’ | 5’-AGAGTACGCTGGCTGAGAGA-3’ |
| II *Ocln* | 5’- ACTCCTCCAATGGCAAAGTG -3’ | 5’- CCCCACCTGTCGTGTAGTCT -3’ |
| *Mbp*^2^ | 5’-AGAGTCCGACGAGCTTCAGA-3’ | 5’-CAGGTACTTGGATCGCTGTG-3’ |
| *Pmp22*^3^ | 5’-GGGATCCTGTTCCTGCACAT-3’ | 5’-TGCCAGAGATCAGTCGTGTGT-3’ |
| *Sox10*^4^ | 5’-GCCACGAGGTAATGTCCAACA-3’ | 5’-TGGTCCAGCTCAGTCACATCA-3’ |
| *Krox20*^4^ | 5’-GCCCCTTTGACCAGATGAAC-3’ | 5’-GGAGAATTTGCCCATGTAAGTG-3’ |
| Taqman Probes Gene | **Taqman assay ID** | **Reference sequence** |
| *Shh*^5^ | Mm00436528_m1 | NM_009170.3 |

^1^murine ribosomal housekeeping gene, ^2^myelin basic protein gene, ^3^peripheral myelin protein gene, ^4^Schwann cell differentiation transcription factors**,** ^5^sonic hedgehog, morphogen

#### Supplemental Table 6. List of primary antibodies used in IHC and Western blotting

| Antibody | Host | Dilution | Application | Source | Cat. No. |
| --- | --- | --- | --- | --- | --- |
| Claudin-1 | Rabbit | 1:100 | IHC | Thermo Fischer | 51-9000 |
| Claudin-5-  -Alexa Fluor 488 | Mouse | 1:100 | IHC | Thermo Fischer | 352588 |
| Claudin-12 | Rabbit | 1:20 | IHC, WB | IBL | 18801 |
| Claudin-19 | Rabbit | 1:100 | IHC | Prof. Hou, St. Louis, USA | (Hou *et al.*, 2009) |
| ZO-1-Alexa Fluor 488 | Mouse | 1:100 | IHC | Thermo Fischer | 339188 |
| CGRP | Rabbit | 1:100 | IHC | ImmunoStar | AIBN 617907 |
| NF200 | Chicken | 1:100 | IHC | Merck | AB5539 |
| IB4-FITC | -- | 10 ug/ml | IHC | Sigma-Aldrich | L2895 |
| SHH^1^ | Mouse | 1:100 | WB | DSHB | AB-2188307 |
| Pan NaCh^2^ | Mouse | 1:100 | IHC | Sigma-Aldrich | S8809 |
| S100b^3^ | Mouse | 1:100 | IHC | Sigma-Aldrich | AMAb91038 |
| β-actin | Mouse | 1:5000 | WB | Sigma-Aldrich | A3854 |
| Occldin | Mouse | 1:100 | IHC | Thermo Fischer | 33-1500 |

^1^ Sonic hedghog, ^2^ pan sodium channel marker, ^3^ Schwann cell marker, Western Blot (WB), immunhistochemistry (IHC)

#### Supplemental Table 7. Quantitation of the tight junction network (Figure 4F).

|  | **WT** | ***Cldn12*-KO** | **p-value** |
| --- | --- | --- | --- |
| **PF associated** |  |  |  |
| **Strand incidence** (%) | 100 (26/26) | 100 (22/22) | --- |
| **Groove incidence** (%) | 0 (26/26) | 0 (22/22) | --- |
| **Particle density*** (n = 22) | 5.76 ± 0.58 | 7.62 ± 0.66 | p < 0.001 |
| **Mesh length** (nm, n = 19) | 412 ± 61 | 638 ± 152 | p < 0.001 |
| **Mesh diameter** (nm, n = 20) | 52.4 ± 17.5 | 26.3 ± 5.6 | p < 0.001 |
| **EF associated** |  |  |  |
| **Strand incidence** (%) | 100 (21/21) | 0 (0/18) | --- |
| **Groove incidence** (%) | 68 (13/19) | 100 (19/19) | --- |
| **Particle density*** (n = 15) | 4.27 ± 0.68 | 2.84 ± 0.96 | p < 0.001 |
| **Mesh length** (nm, n = 14) | 397 ± 73 | 525 ± 83 | p < 0.001 |
| **Mesh diameter** (nm, n = 14) | 71.4 ± 40.3 | 38.0 ± 26.7 | p < 0.01 |

All data are shown as mean ± SEM. Two-tailed student’s t-test was used to determine the statistical significance. In addition, the EF groove intensity rose from 27.6 ± 24.9% in WT to 76.3 ± 29.4% in *Cldn12*-KO, *P* < 0.01. Particle density*: per 100 nm strand and groove, respectively.

**Supplemental Methods**

The following mouse phenotyping and resulting **Supplemental Fig S3** were provided by Mouse Clinical Institute, Illkirch/France (https://www.mousephenotype.org/impress).

***Mouse phenotyping:*** Nine male *Cldn12*-KO and 9 WT littermates were analyzed throughout the IMPC core phenotyping pipeline and the IMPRESS procedures (<https://www.mousephenotype.org/impress>) covering the main physiological and behavior functions. All mice were fed a standard chow diet (D04-Safe) and allowed to acclimatize in our laboratory animal facility.

*Rotarod test*: The ability of an animal to maintain balance on a rotating rod (Bioseb, Chaville, France) was tested as index of motor coordination performance. Mice were given three testing trials during which the rotation speed accelerated from 4 to 40 rpm in 5 min. Average latency time until animals fell off the rod were recorded. Trials were separated by a 5‐10 min interval.

*Grip test*: Maximal muscle strength (g) was measured using an isometric dynamometer connected to a grid (Bioseb). Mice were allowed to grip the grid with all their paws. Then, they were pulled backwards until they released it. Each mouse was submitted to 3 consecutive trials immediately after the modified SHIRPA procedure. The maximal strength developed by the mouse before releasing the grid was recorded and the average value of the three trials adjusted to body weight.

*Hot plate test*: Mice were placed into a glass cylinder on a hot plate adjusted to 52°C (Bioseb) and the latency of the first pain reaction of any hindlimb (e.g. licking, flinches) was recorded, with a maximum of 30 s.

*Shock threshold test*: Mice were placed in a fear‐conditioning chamber and allowed to habituate for 30 s. An electrical foot‐shock was then manually applied for 1 s and behavioral responses noted. Shock levels began at 0.05 mA, and increased in 0.05 mA steps with 30 s interval between the shocks until both flinch (any detectable response) and vocalization were induced. After this point, shocks were increased in 0.1 mA steps until a jump (the mouse flinches such that the two hind paws leave the ground) was induced. A 1 mA cut‐off was employed in this test.

*Auditory brainstem responses (ABR)*: Auditory brainstem response test determines hearing sensitivity using evoked potential recordings in anaesthetized mice. The ABRs were recorded using an electrophysiological station composed of different items from Tucker‐Davis Technologies (Alachua, FL, USA). Mice were anesthetized with a mixture of Ketamine‐Xylazine and then placed in a sound‐attenuating chamber facing a loud‐speaker and recording electrodes appropriately placed on the skull. ABRs were recorded to different acoustic stimuli. ABRs were first noted to clicks (white noise) (10 μs duration, positive transient) presented from 0‐85 dB SPL in 5dB steps, presented 256 times at 42.6/s. ABRs were then recorded to the following frequencies and intensities of stimuli; 6 kHz (20‐85 dB SPL), 12 kHz (0‐70 dB SPL), 18 kHz (0‐70 dB SPL), 24 kHz (10‐70 dB SPL) and 30 kHz (20‐85 dB SPL), presented in 5 dB intervals. (Tone pips are 5 ms in duration, with a 1 ms rise/fall time, presented 256 times at 42.6/s). At the end of analysis, a final recording for clicks was performed. After recovery from anesthesia, mice were returned to their home cages.

*Auditory Startle Reflex Reactivity and Pre‐Pulse Inhibition*: Acoustic startle reactivity and pre‐pulse inhibition of startle were assessed in a single session using standard startle chambers (SR‐Lab Startle Response System, San Diego Instruments, USA). Ten different trial type were used: acoustic startle pulse alone (110‐db), eight different prepulse trials in which either 70, 75, 85 or 90‐dB stimuli were offered alone or preceded the pulse, and finally one trial (NOSTIM) in which only the background noise (65 dB) was presented to measure the baseline movement in the Plexiglas cylinder. In the startle pulse or prepulse alone trials, the startle reactivity was analyzed and in the prepulse plus startle trials the amount of PPI was measured and expressed as percentage of the basal startle response.

*The pavlovian fear conditioning*: Polymodal operant chambers (Coulbourn Instruments, Allentown, PA, USA) were used. Each chamber (18.5 x 18 x 21.5 cm) consisted of aluminum side walls and plexiglas rear and front (the door) walls. A loudspeaker and a bright light were cues sources during conditioning and cue‐testing. The general activity of animals was recorded through the infrared cell placed at the ceiling of the chambers connected to a PC computer using the Graphic State software (Coulbourn). For conditioning, mice were allowed to acclimate for 4 min, then a light/tone (10 kHz) CS was presented for 20 s and co‐terminated by a mild (1 s, 0.4 mA) foot shock (US). Mice were returned to their home cages 2 min later. Testing was performed 24 h following conditioning session. Testing for the context was accomplished in the morning. Mice were placed back into the same chamber that was used for the conditioning and allowed to explore for 6 min without presentation of the light/auditory CS. Testing for the cue was performed in the afternoon (about 5 h after the context testing). The contextual environment of the chambers was changed (wall color, odor and floor texture). Mice were placed in the new chamber and allowed to habituate for 2 min then presented with light/auditory cues for 2 min. This sequence was repeated once again. At the end of testing, animals were returned to their home cages.

*Vision exploration*: Slit lamp examination of the anterior segment of the eye (cornea, iris, lens) did not reveal any difference between *Cldn12*-KO male mice and WT littermates (9 mice/18 eyes for each genotype). Minor defects, as mild suture cataracts or spot cataracts (some of them linked to focal persistent hyperplastic primary vitreous) were seen in both genotypes, as classically for a C57Bl/6N background. Central retina was monitored on a diameter of 1.4 mm, on both eyes of WT and *Cldn12*-KO mice. For each eye, the thickness of the following elements were measured, at a distance of ~0.5 mm of the optic nerve, chosen as representative (values are mean ± SD): Total retinal (from the neuronal fiber layer to the pigmentary epithelium), photoreceptor segments, outer Nuclear Layer (ONL, formed by the photoreceptor nuclei), inner Nuclear Layer (INL, formed by the nuclei of bipolar and amacrine cells), inner Plexiform Layer (IPL, formed by the axons of bipolar cells and the dendrites of amacrine and ganglion cells).

***Genotyping:*** Ear punch biopsies (1-2 mm^2^) from *Cldn12*-KO and WT mice were obtained. The tissue samples were incubated in mixed lysis buffer (5 µl proteinase K with 50 µl lysis buffer) at 55 °C in a heating incubator overnight. Lysed tissue solution (1 µl) containing DNA sample was mixed with corresponding master mixes. PCR was used to genotype KO and WT mice. PCR amplification was performed in 2720 Thermal Cycler (Thermo Fisher Scientific). The PCR products were detected by agarose gel electrophoresis.

***Histology and histomorphometric evaluation:*** SN tissues of naïve KO and WT mice were collected and fixed with 2.5% glutaraldehyde. 2% osmium tetroxide (Sigma-Aldrich, St. Louis, USA) was used for post-fixation. Post-fixed nerves were embedded in the Spurr's resin (plastic) before cutting. Semi-thin sections (0.5 µm) were prepared and stained with azure-methylene blue. Subsequently, the morphology of nerve fibers and myelin sheaths were assessed to detect pathological findings by light microscopy using the microscope KEYENCE BZ-9000 immunofluorescence microscopy (Osaka, Japan). Axon diameter in relation to the total diameter of the nerve fiber (g-ratio) and the number of myelinated fibers per area were measured and analysed. Fibers with undulated myelin were defined as fibers with >4 undulations on myelin as previously observed in aging mice (Yuan *et al.*, 2018). The number of fibers per area was used to calculate the axon density of single nerves.

***Real-time qPCR:*** Whole SN were microsurgically separated into epiperineurium (EPN) the desheathed SN (dSN). Total RNA was extracted from the DRG, dSN and EPN using miRNeasy Micro kit (Qiagen, Düsseldorf, Germany) following the manufacturer’s protocol (Sauer *et al.*, 2014; Sauer *et al.*, 2017; Reinhold *et al.*, 2019). RNA concentration was determined with the peqlab NanoDrop ND 2000 spectrophotometer (Fisher Scientific, Schwerte, Germany). cDNA was synthesized from 1 µg of total RNA with high capacity cDNA reverse transcriptase Kits (with random primer and RNAse inhibitor; Invitrogen, Carlsbad, CA, USA) according to the manufacturer’s instruction. A 10x dilution of cDNA was used in quantitative reverse transcriptase–PCR experiments. All the PCR measurements were performed in triplicates with StepOnePlus Real-Time PCR System (Thermo Fisher Scientific) and PowerUp^TM^ SYBR^®^Green. Ct values were averaged for relative quantification. mRNA expression was normalized to the mouse ribosomal housekeeping gene S18 and displayed as fold change expression calculated by the 2^-ΔΔCT^ method (Yang *et al.*, 2016; Reinhold *et al.*, 2019). We listed primer sequences in the **Supplemental Table 5.** Primers were designed using Primer3 software after verification by using FASTA searches.

***Western blotting:*** Proteins were isolated and pooled from the dSN, EPN and DRG and homogenized by TissueLyser (QIAGEN) after freeze-thawing in Trizol (Invitrogen). The lysed protein (20 µg) was mixed with SDS containing buffer containing 1X Laemmli buffer and 6% ß-mercaptoethanol. Then, they were denatured at 55 °C for 2 min 30 sec. Western blotting was performed after loading the denatured proteins into each lane on a 4%–12% Tris-glycine gel (Invitrogen). We used 12% SDS polyacrylamide gels to fractionate total protein from samples and subsequently blotted the gels overnight onto Protran^®^ Nitrocellulose membrane (Sigma, GE10600001). Membranes were blocked with 5% non-fat milk powder (AppliChem, Darmstadt, Germany) at RT for 1 h. Subsequently, membranes were incubated with primary antibodies at 4 °C overnight. The mouse monoclonal anti-β-actin HRP Ab (1:2500; Invitrogen) was used as the loading control. Primary antibodies and their dilutions are listed in the **Supplemental Table 6.** Blots were further incubated with the horseradish peroxidase (HRP)-conjugated secondary antibodies diluted with 1% BSA in PBS. Labeled proteins were detected by chemiluminescence (ECL solution, Pierce) and densitometry (FluorChem FC2 Imaging systems, Multilmage II; Alpha Innotech). Gray scale images and bands’ intensity were analyzed using the open source software ImageJ (plot analyzed tool), and were calculated as expression relative to β-actin protein expression.

***Immunofluorescence:*** DRG and SN tissues were dissected, embedded into Tissue–Tek (O.C.T. TM compound containing), cryo-sectioned at 10 µm and mounted on SuperFrost*/plus microscope slides. Slides were baked for 15 min at 37 °C before storage at –20 °C. Sections were incubated for 7 min with 0.5% Triton X-100 (Sigma-Aldrich, St Louis, MO, USA) in PBS after acetone fixation for 15 min at -20°C. We used 10% donkey-serum to block non-specific binding in tissues for 90 min at RT. Then, tissue sections were incubated with primary antibodies overnight at 4 °C (**Supplemental Table 6**) followed by the corresponding fluorescence-conjugated secondary antibodies (1:1000 in PBS) for 1 h at RT. We washed samples with PBS and ddH_2_O before mounting with Vectashield^TM^ Mounting Medium (Vector Labs, Burlingame, CA, USA).

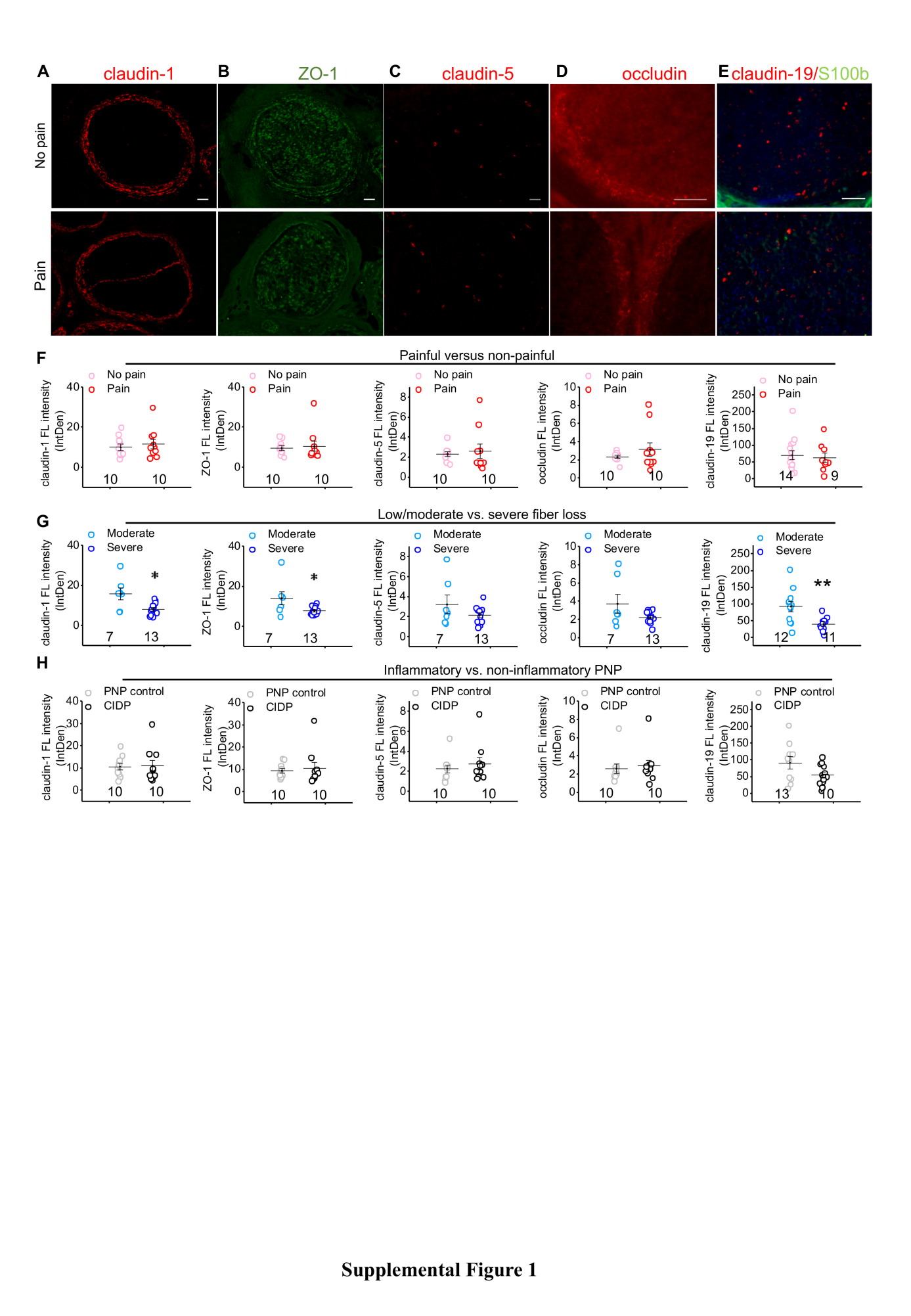

**Supplemental Figure S1.** **Reduced immunoreactivity of claudin-1, -19 and ZO-1 but not claudin-5 or occludin in patient with chronic inflammatory demyelinating polyneuropathy (CIPD) or non-inflammatory PNP with severe fiber loss.** Sections from sural nerves biopsies of patients with either noninflammatory PNP or CIDP were immunolabeled. Patients’ characteristics are described in **Supplemental Table 1** (20 patients; claudin-1, -5, occludin and ZO-1) and **Supplemental Table 2, 3** (23 patients; claudin-19). Representative immunostainings staining for claudin-1 (**A**), ZO-1 (**B**), claudin-5 (**C**), occludin (**D**) and claudin-19 co-stained with the Schwann cell marker S100b (green) (**E**) in the sural nerve from patients with painful or nonpainful noninflammatory PNP or CIDP (scale bar: 50 µm). (**F-H**) Immunofluorescent stainings were quantified: claudin-1 was analyzed in the perineurium only, ZO-1 and occludin in the whole nerve, claudin-5 and claudin-19 in the nerve without the perineurium (endoneurium). Data were compared in patients with noninflammatory PNP or CIDP with or without pain (**F**), with low/moderate fiber loss compared to with severe fiber loss (**G**) and from CIDP patients compared to noninflammatory PNP patients (**H**). *p < 0.05, **p < 0.01 two-tailed student’s t-test. The number of patients in each group are displayed below the dots. Data are shown as mean ± SEM.

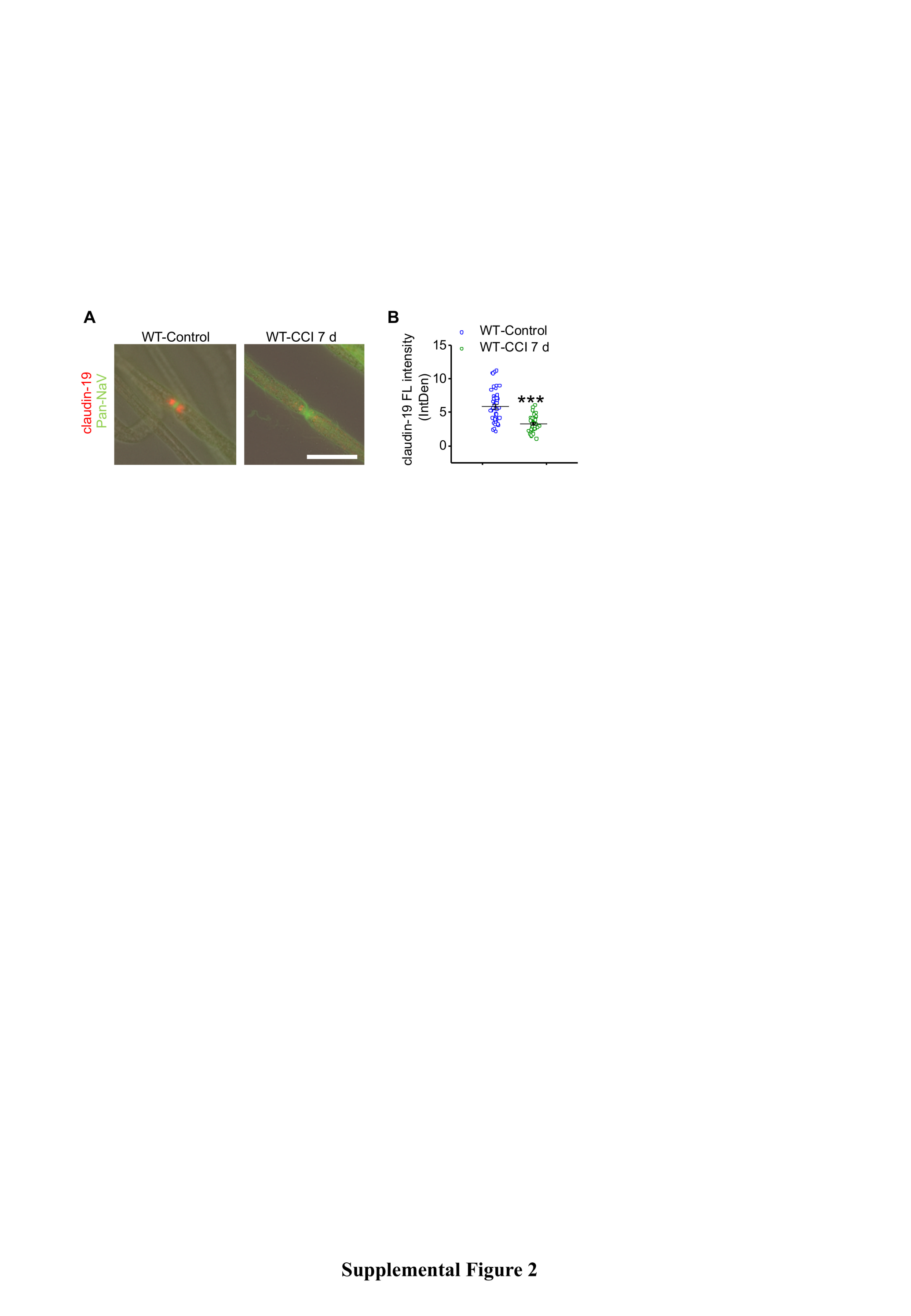

**Supplemental Figure S2.** **Loss of claudin-19 in the paranode after chronic constriction injury** (**CCI)**. WT mice underwent CCI for 7 d. (**A**) Teased fibers were stained for claudin-9 (red) and nodes identified by Pan-NaV Ab (green). (**B**) Claudin-19-IR was quantified in the paranodal region untreated control and CCI mice. (***p < 0.001, one-tailed Student’s t-test; n = 40-50 fibers from six mice per group, scale bar: 20 µm).

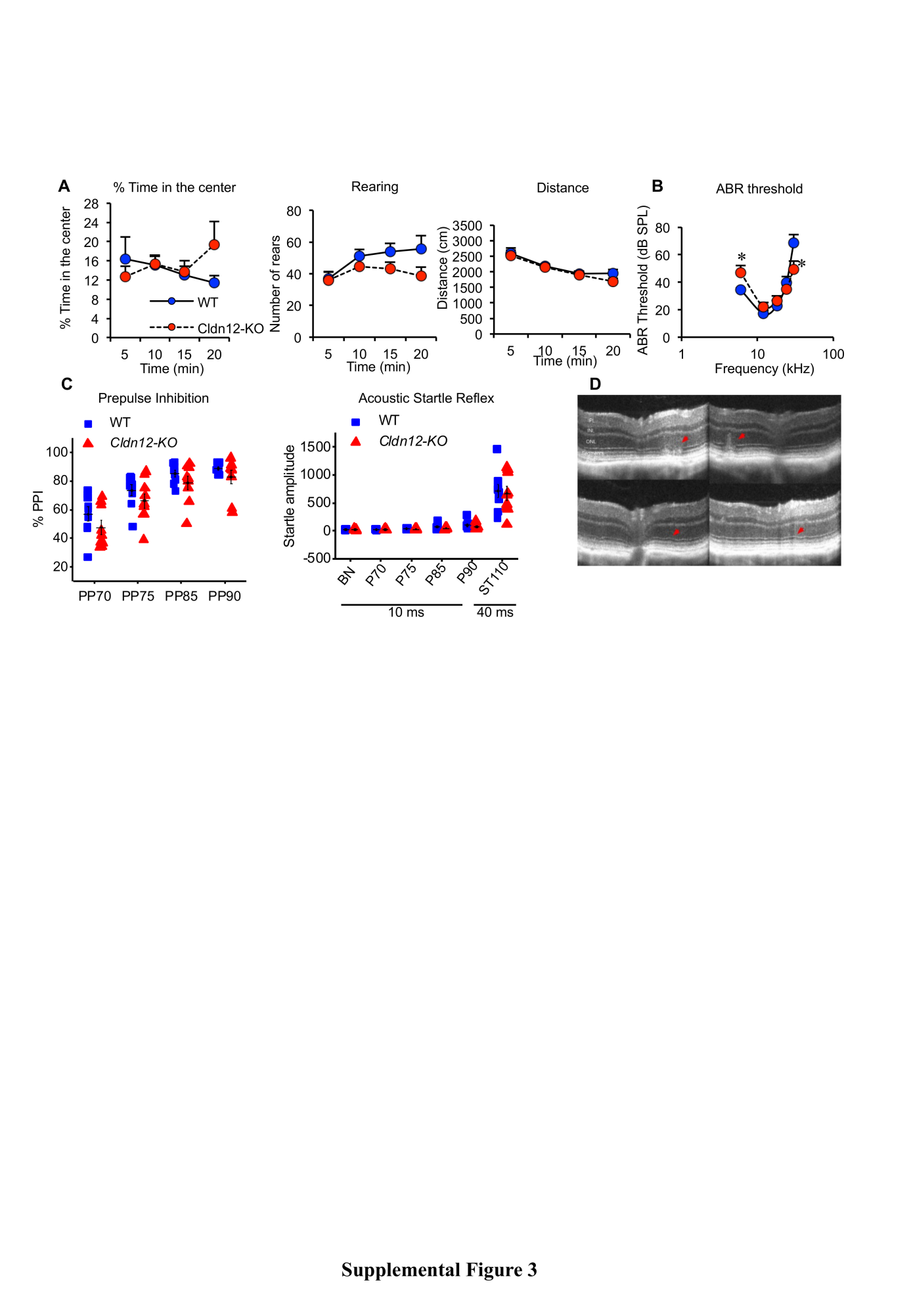

**Supplemental Figure S3. No differences in anxiety-related behavior, sensorimotor gating and auditory function of *Cldn12*-KO mice.** Male WT and *Cldn12*-KO mice were subject to a panel of neurological test for anxiety-related behavior, sensorimotor gating as a model for schizophrenia as well as visual and auditory function. (**A**) In the open-field test, locomotor activity and rears were comparable between WT and KO male mice. Exploration of the central part of the arena did not have significant difference between these two genotypes, suggesting the anxiety-related behavior was not affected in mutants. (**B**) The consequence of *Cldn12* deletion on acoustic startle and pre-pulse inhibition (PPI) of startle reflex were also evaluated. The startle reactivity was comparable between WT and KO males for all the acoustic stimuli used including the startling pulse. The level of PPI was also comparable between genotypes. Comparable result from two startle tests indicated that *Cldn12* deletion did not drag in the reflexive contraction of the skeletal musculature in response to strong exteroceptive stimuli. (**C**) Specific evaluation of the auditory system using auditory brain stem responses did not reveal any alteration in mutants; the audiograms were comparable between genotypes. (**D**) Retinas of KO mice were slightly but significantly (p = 0.00018) thinner than those from their WT littermates. This difference came from a thinner inner retina (INL + IPL), the IPL being significantly thinner in mutant mice (p = 0.013), while the outer retinas (photoreceptors) were comparable in between genotypes. Due to the C57Bl/6N background, classical signs of the rd8-associated retinal degeneration were observed, from large spots of dysplasia (top row) to small ripples in the external limiting membrane (bottom row), with comparable severity and distribution between WT and mutant mice.

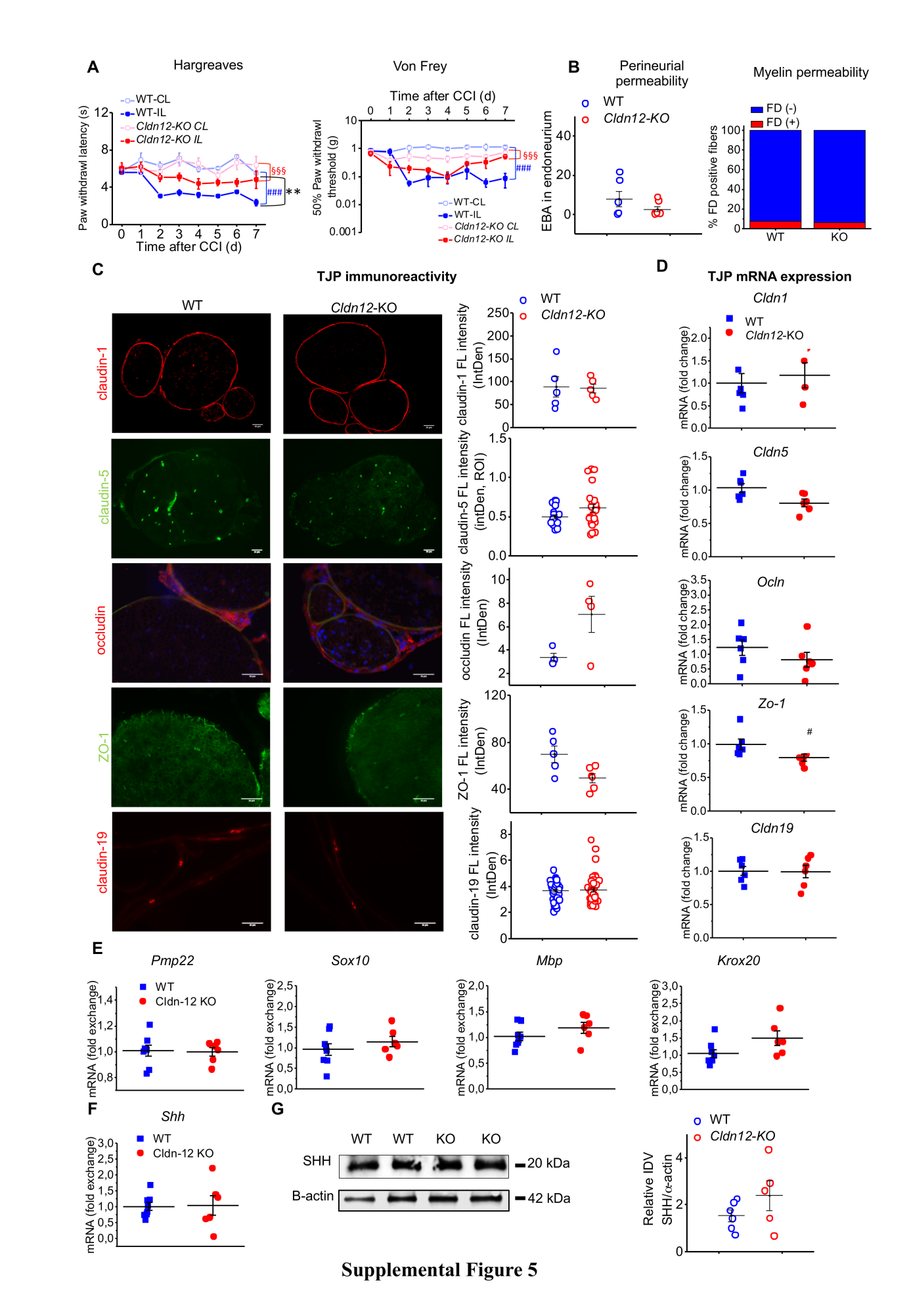

**Supplemental Figure S4. No significant difference in thermal and mechanical nociceptive responses and barrier permeability are observed in female *Cldn12*-KO mice**. (**A**) Thermal nociceptive thresholds (Hargreaves) and mechanical nociceptive thresholds (von Frey) were determined in female WT and KO mice following 1-7 d after chronic constriction injury (CCI). (n = 6, Two-way RM-ANOVA, Bonferroni’s post hoc test. **p < 0.001 IL *Cldn12*-KO versus IL in WT in the Hargreaves test only. §§§ p < 0.001 IL versus CL *Cldn*12-KO, ### p < 0.001 IL versus CL WT). Data are shown as mean ± SEM. IL: ipsilateral, CL: contralateral. (**B**) Perineurial permeability was assessed and quantified after plasma albumin associated Evans Blue (EBA, 68 kDa) incubation in the sciatic nerve (SN). Myelin permeability was assessed by as well as the number of FITC dextran (FD; 70 kDa) positive and negative fibers from WT and *Cldn12*-KO mice SN. (**C, D**) Tight junction protein immunoreactivity (left) and its quantification was measured in the regions of interest (ROI) (middle). The mRNA expression of *Cldn1*, *Cldn5*, *Cldn19*, *Ocln* and ZO-1 (*Tjp1*) (right) were quantified in the desheathed sciatic nerve (dSN: *Cldn5*, *Cldn19*, *Ocln* and *Tjp1*) or epiperineurium (EPN: *Cldn1*). (**E**) From left to right: mRNA quantification of Schwann cell markers or differentiation markers *Pmp22*, *Sox10*, *Mbp* and *Krox20* in the SN. (**F**) Analysis of sonic hedgehog (*Shh*) mRNA expression, representative Images of Western blot and quantitative result of SHH expression from the SN of WT and *Cldn12*-KO mice (B-F: n = 3-6/group, p > 0.05, two-tailed Student’s t-test).

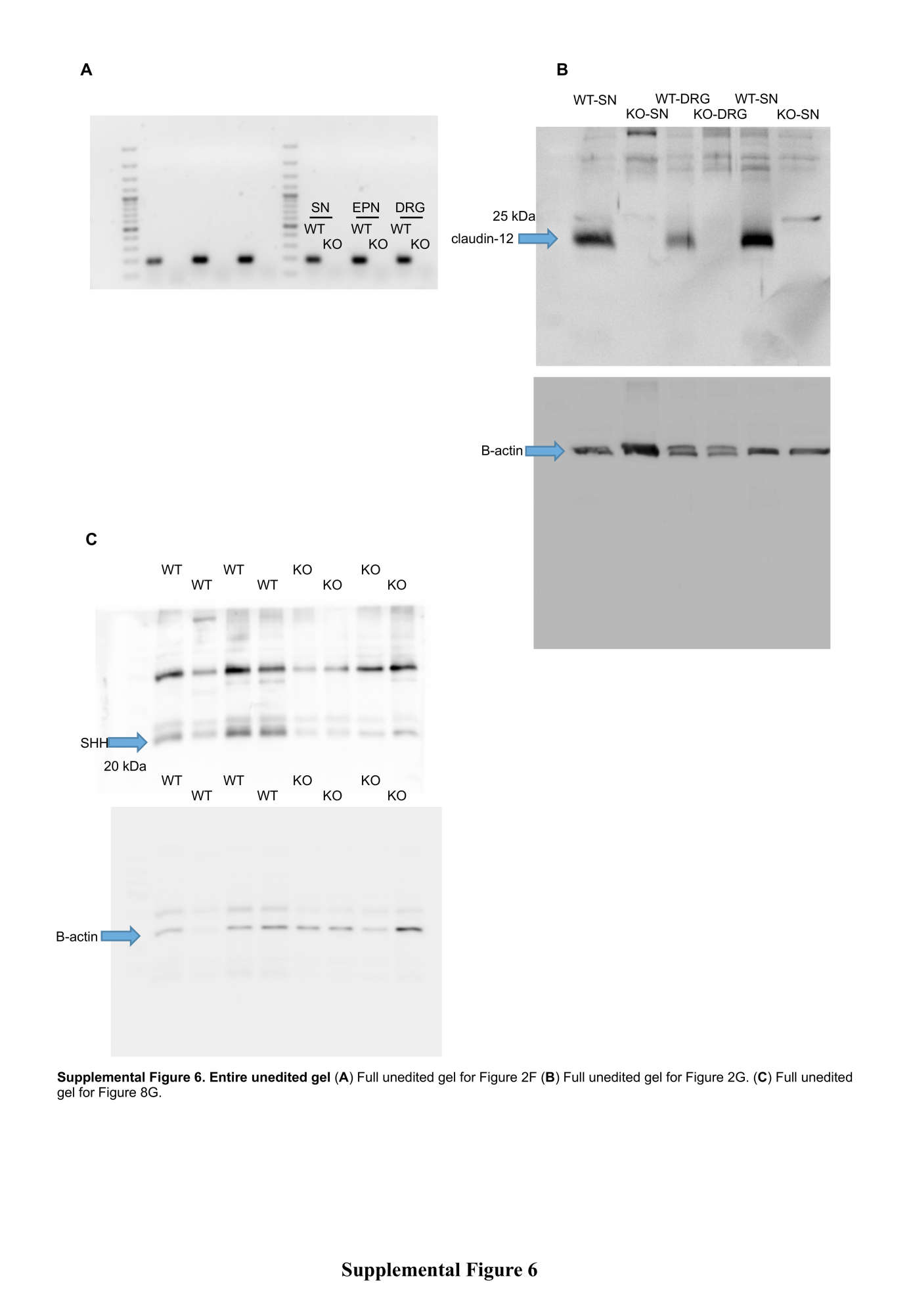

**Supplemental Figure S5. Entire unedited gel** (**A**) Full unedited gel for **Fig 2E**. (**B**) Full unedited gel for **Fig 2F**. (**C**) Full unedited gel for **Fig 8G**.
